# A Multimodal Framework to Uncover Drug-Responsive Subpopulations in Triple-Negative Breast Cancer

**DOI:** 10.1101/2025.02.14.638274

**Authors:** Yue Wang, Santiago Haase, Austin Whitman, Adriana Beltran, Philip M. Spanheimer, Elizabeth Brunk

**Author notes:** Correspondence should be addressed to: Elizabeth Brunk; Philip Spanheimer.

## Abstract

Understanding how individual cancer cells adapt to drug treatment is a fundamental challenge limiting precision medicine cancer therapy strategies. While single-cell technologies have advanced our understanding of cellular heterogeneity, efforts to connect the behavior of individual cells to broader tumor drug responses and uncover global trends across diverse systems remain limited. There is a growing availability of single-cell and bulk omics data, but a lack of centralized tools and repositories makes it difficult to study drug response globally, especially at the level of single-cell adaptation. To address this, we present a multimodal framework that integrates bulk and single-cell treated and untreated transcriptomics data to identify drug responsive cell populations in triple-negative breast cancer (TNBC). Our framework leverages population-scale bulk transcriptomics data from TNBC samples to define seven main “identities”, each representing unique combinations of biologically relevant genes. These identities are dynamic and trackable, allowing us to map them onto single cells and uncover global patterns of how cell populations respond to drug treatment. Unlike static classifications, this approach captures the evolving nature of cellular states, revealing that a select few identities dominate and drive population-level responses during treatment. Crucially, our ability to decode these trends through the inherent noise of single-cell data provides a clearer picture of how heterogeneous cell populations adapt to therapy. By identifying the dominant identities and their dynamics, we can better predict how entire tumors respond to treatment. This insight is essential for designing precise combination therapies tailored to the unique heterogeneity of patient tumors, addressing the single-cell variations that ultimately determine therapeutic outcomes.

---

How cancer cells adapt to drug treatment is one of the most pressing challenges in cancer research. Understanding how tumor cells respond, adapt, and resist therapies is essential for developing strategies that target these distinct populations within tumors. Within a single tumor, cells can respond to therapy in dramatically different ways^1^: some are eliminated, while others adapt, survive, and drive resistance^2,3^. This response heterogeneity undermines even the most promising therapies^4–6^, making it critical to identify which subpopulations of cells are responsible for emerging resistance.

In recent years, the explosion of omics technologies has produced an unprecedented wealth of data. We now have access to transcriptomic profiles spanning diverse cancer types, experimental conditions, and molecular modalities at bulk and single-cell levels. This includes untreated and drug-treated samples, offering a rare opportunity to explore how cancer cells adapt at the molecular level. However, the sheer volume and complexity of these datasets present significant challenges. Distinguishing meaningful biological signals from noise amid thousands of single cells remains a daunting computational task^7–10^. In the context of clinical research, this challenge extends to aggregating data into therapeutically meaningful patterns while maintaining the granularity needed to characterize rare or low-abundance cell populations. Despite numerous efforts to address this complexity^11–18^, current approaches often focus on predicting drug responses and lack the corresponding single-cell omics data in the drug treatment setting, leaving many adaptive mechanisms unresolved and limiting our ability to uncover global trends in drug response.

Efforts to integrate datasets across untreated and treated conditions have been hindered by the lack of unified repositories and comprehensive analytical tools, though longitudinal datasets exist^19–23^. As a result, most studies of drug response focus on deep investigations of individual cell lines or tumor models^24–28^. While these studies offer valuable mechanistic insights, they are often limited in scope and provide little opportunity to identify trends that are broadly applicable across samples in a population. The field urgently needs scalable methods capable of uncovering population-wide patterns—tools that can illuminate how adaptation arises, predict drug responses across diverse cancers, and inform the development of more effective therapies. Moreover, fully leveraging the wealth of available multiomics data requires approaches that bridge bulk and single-cell datasets^29–31^, integrating large-scale trends with single-cell granularity. Addressing these gaps would represent a significant step forward in advancing precision medicine.

In this study, we tackle these challenges by introducing a multimodal framework that integrates bulk and single-cell transcriptomics data to uncover population-scale trends in drug response. We apply this framework to triple-negative breast cancer (TNBC), a highly aggressive subtype of breast cancer that lacks effective targeted therapies and is marked by variable treatment responses^32,33^. By harmonizing bulk and single-cell datasets from untreated and drug-treated conditions, our framework identifies transcriptionally distinct subpopulations with specific molecular “identities” within TNBC cell lines and tumors. By grouping thousands of cells into biologically relevant identities, this approach enables the identification and characterization of subpopulations of cells that may be driving treatment-induced changes, offering a critical lens into how tumors adapt and evolve under therapeutic pressure.

By capturing population-scale patterns and linking them to single-cell behaviors, our framework addresses a critical unmet need in cancer research. It enables the identification of nuanced, biologically meaningful shifts in cellular identities that drive adaptation and resistance, ensuring critical signals are not lost in complex data. Our analysis revealed striking adaptive signatures, with the largest transcriptional shifts often associated with a small number of highly adaptive subpopulations of cells characterized by key identities. As a proof of principle, we applied our framework to our TNBC organoid model, collecting single-cell multiomics data from untreated and treated samples following treatment with epigenetic remodeling drugs. Analysis of this model validated that the predicted identity shifts correspond to distinct epigenetic regulatory mechanisms, highlighting the biological relevance of our approach. Beyond TNBC, this approach has the potential to uncover actionable molecular vulnerabilities across cancer types, paving the way for new precision medicine strategies to outpace resistance and improve patient outcomes.

## Results and Discussion

### Harmonizing Bulk and Single-Cell Transcriptomics for Defining Subpopulation Identities

To establish a comprehensive framework for defining subpopulation identities, we curated publicly available datasets focusing on triple-negative breast cancer (TNBC). We identified 30 cell lines annotated as TNBC from the Cancer Cell Line Encyclopedia (CCLE) and Dependency Map (DepMap), which included 76 transcriptomic datasets, 30 bulk and 46 single cell datasets^34–36^. For the single cell data, five have corresponding drug-treatment datasets^37–39^. These datasets offered a robust foundation for exploring the molecular diversity of TNBC, encompassing both untreated and treated conditions. The availability of multiomics data allowed us to harmonize bulk and single-cell transcriptomics, enabling detailed analysis of cell subpopulation variability (**Fig. 1A**).

**Figure 1.**
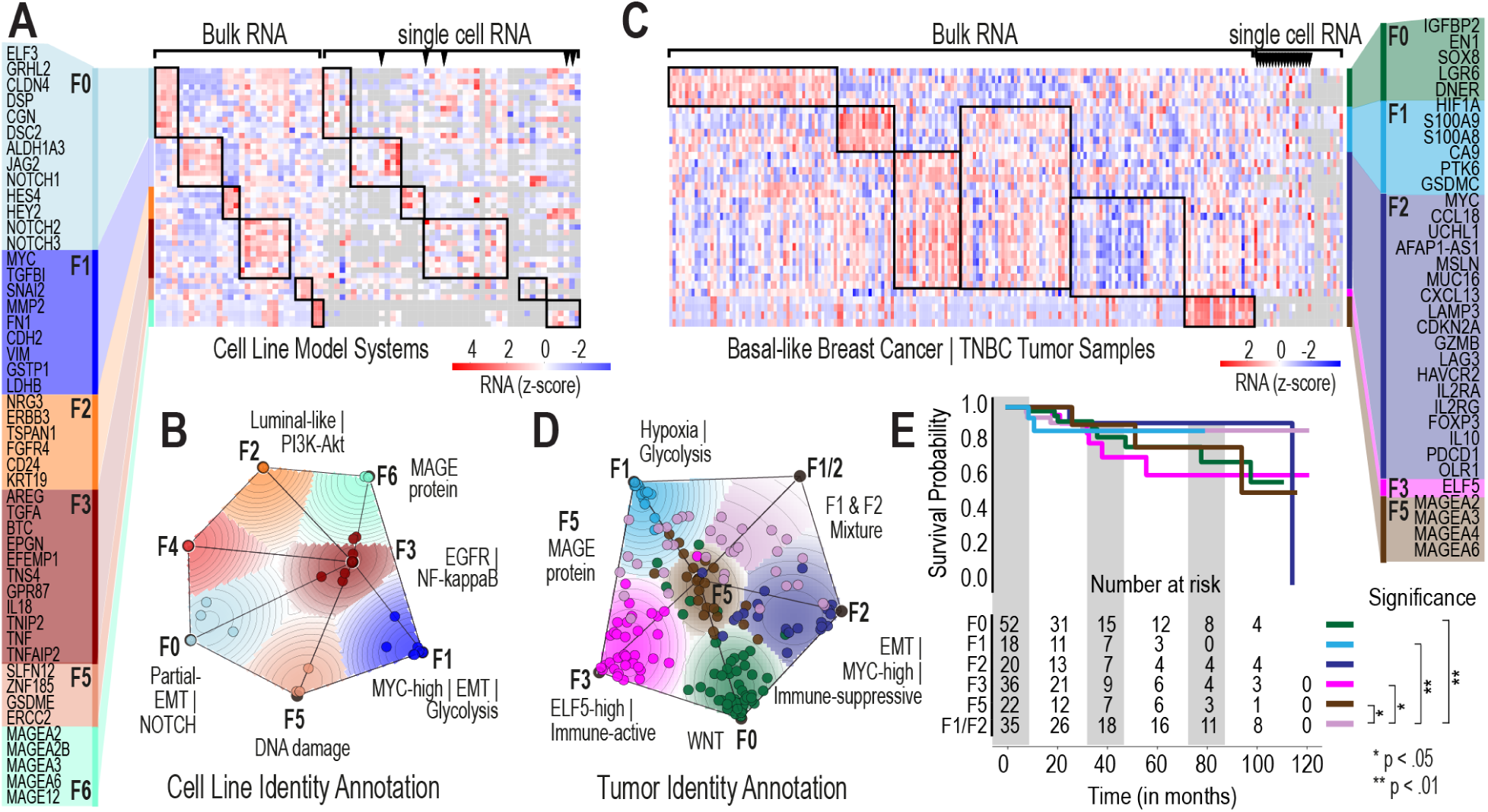
Gene Expression Identity Landscapes for TNBC Cell Lines and Tumor Samples. **1A.** Heatmap illustrating the overall transcriptomic data resource for CCLE TNBC cell lines. Each column represents a cell line, and each row corresponds to the expression of genes defining specific identities. Identities and their representative gene groups are annotated on the left side of the heatmap. Samples associated with the same identity are outlined with black boxes, while single-cell RNA-seq data with corresponding drug treatment conditions are indicated along the top edge of the heatmap. Black triangles indicate drug treated samples. **1B.** Transcriptomic identity map for 30 TNBC cell lines, highlighting the identity groupings derived from CCLE data. **1C.** Heatmap of the transcriptomic data resource for basal-like breast cancer and TNBC tumor samples. Each column represents a tumor sample, and each row corresponds to the expression of genes defining specific identities. Identities and their associated gene groups are annotated on the right side of the heatmap. Samples grouped by the same identity are outlined with black boxes, and patient single-cell RNA-seq data with corresponding drug treatment conditions are marked along the top edge of the heatmap. **1D.** Transcriptomic identity map for 185 TCGA basal-like breast tumor samples, highlighting the identity clusters derived from TCGA data. **1E.** Kaplan-Meier survival curves for TCGA basal-like breast tumor samples, stratified by identity. The risk table indicates the number of patients at risk over time for each group, including the F1/F2 Mixture identity, denoted as F1/F2.

To identify the genes that best define distinct subpopulations, we performed a genome-wide analysis to pinpoint those with the highest variance across TNBC cell lines. Given the well-established role of MYC as a global gene regulator impacting TNBC^40–43^, we prioritized significantly varied genes and examined pair-wise relationships between MYC and each gene to identify those that most distinctly separated the TNBC cell lines into discrete clusters. This analysis identified 523 genes with highly divergent expression patterns, providing a basis for separating transcriptionally distinct subpopulations. As expected, these genes highlighted diverse biological pathways, including MYC dysregulation, a hallmark of TNBC biology, as well as other processes critical for understanding adaptive responses to treatment, including NOTCH, PI3K-Akt and EGFR signaling.

Using the most divergent genes, we applied non-negative matrix factorization (NMF) to identify transcriptional groups, or “identities,” among TNBC cell lines. This approach revealed seven subpopulation identities, each characterized by unique gene expression profiles that capture the heterogeneity across 30 TNBC cell lines. These identities were further visualized using a two-dimensional cluster map, which clearly illustrated the distinct groupings of TNBC cell lines (**Fig. 1B**).

To provide biological context, we performed detailed literature searches (**Supplementary Table 1-6**) to annotate each identity, focusing on the most relevant biological features (**Supplementary Information Fig. 1-7**) associated with the defining genes. For example, Identity F2 was enriched for luminal-like genes and PI3K-Akt signaling, while Identity F1 was linked to markers of epithelial-to-mesenchymal transition (EMT) and glycolysis. These annotations illuminated the diverse molecular processes driving subpopulation variability, offering insights into potential therapeutic targets.

We extended our analysis to The Cancer Genome Atlas (TCGA) basal-like breast cancer datasets to evaluate the relevance of these subpopulation identities in patient samples (**Fig. 1C**). Notably, several subpopulations identified in cell lines were conserved in TCGA tumors, suggesting that these transcriptional identities are biologically meaningful and clinically relevant (**Fig. 1D, Extended Data** Fig.1 **and Supplementary Information Fig. 8-11**). For instance, Identities F5 and F1-and-F2-mixture characterized by genes involved in MAGE protein family and high-MYC & hypoxia/glycolysis programs respectively, were consistently observed across both datasets (**Extended Data** Fig.1B).

Importantly, we found that certain subpopulations assigned to the TCGA samples were associated with significant differences in patient survival outcomes. Subpopulation F1-and-F2-mixture, distinguished by both high hypoxia and immune-suppressive genes, exhibited better overall survival compared with F0 subpopulation (p= 0.005), F1 subpopulation (p=0.006), F3 subpopulation (p=0.014) and F5 subpopulation (p=0.02) (**Fig. 1E)**. This observation highlights the potential of subpopulation-based analyses to stratify patients into clinically relevant prognostic groups which could be used to guide therapeutic decision-making linked to drug responsiveness.

Through bulk transcriptomics analysis, we identified transcriptionally distinct subpopulations, referred to as identities, that capture the inherent heterogeneity of TNBC. These identities lay the groundwork for understanding the molecular processes underlying variability in TNBC outcomes and demonstrate their clinical relevance, presenting new opportunities for precision medicine in this aggressive cancer subtype. Additionally, these findings establish a baseline understanding of TNBC cell identities in treatment-naive conditions, providing a crucial foundation for investigating drug response trajectories under therapeutic pressure.

### Integrating Multimodal Data to Define and Validate Subpopulation Identities

We used harmonized multimodal data to validate the transcriptionally distinct subpopulation identities defined in TNBC cell lines. This harmonized dataset integrates bulk RNA sequencing, CRISPR loss-of-function screens, and drug sensitivity data from repositories such as the Cancer Therapeutics Response Portal (CTRP)^44,45^ and the Genomics of Drug Sensitivity in Cancer (GDSC)^46–48^ (**Fig. 2A**). By combining these distinct phenotypic datasets, we aimed to independently validate and characterize the biological relevance of the TNBC subpopulation identities identified through NMF.

**Figure 2.**
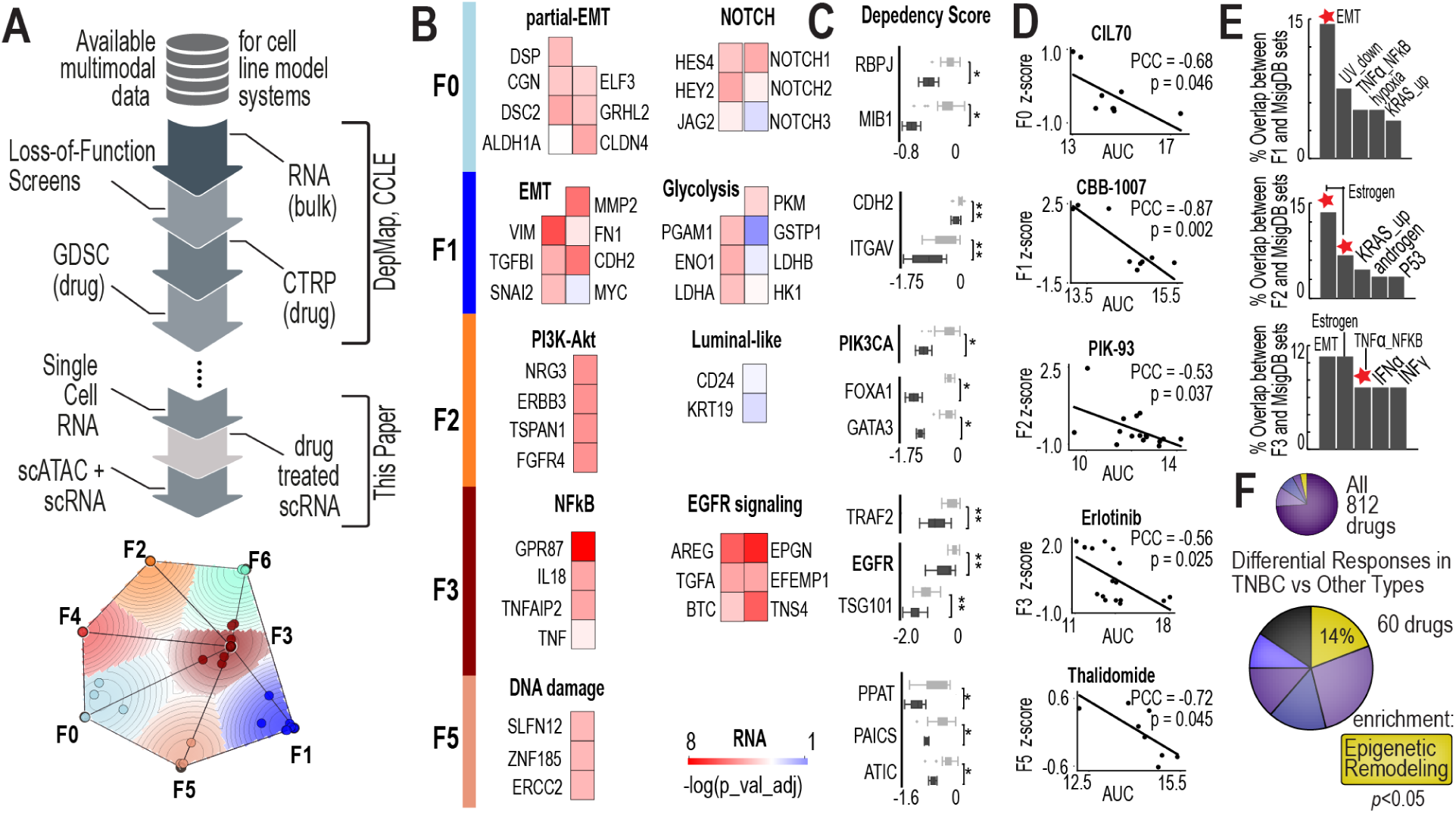
Multi-Omics Evidence Supporting the Biological Relevance of TNBC Cell Line Identities. **2A.** The workflow for annotating TNBC cell line identities, providing an overview of the methodology used to assign biologically meaningful features to these identities. **2B.** Gene expression evidence supporting the biology of TNBC cell line identities. The heatmap presents the adjusted p-values from Wilcoxon rank-sum tests comparing gene expression levels in cell lines belonging to a specific identity versus all others. Genes representative of each identity show significantly higher expression within the associated samples, confirming their relevance to the defined biological features. **2C.** CRISPR dependency data validates the biological characteristics of each TNBC cell line identity. Dependency scores are shown for cell lines belonging to each identity compared to those outside it. Dark gray box plots represent cell lines associated with the identity, while light gray box plots represent others. For example, RBPJ and MIB1 highlight NOTCH signaling in the F0 identity; CDH2 and ITGAV emphasize EMT features in F1; PIK3CA, FOXA1, and GATA3 define PI3K-Akt signaling and luminal-like traits in F2; and TRAF2, EGFR, and TSG101 highlight NF-κB and EGFR signaling in F3. **2D.** Linking drug sensitivity to gene expression identities within TNBC cell lines. Correlations between identity levels and drug sensitivity are depicted with Pearson correlation coefficients (PCCs) and p-values. Key examples include drugs such as CIL70 (ferroptosis inducer), CBB-1007 (histone demethylase inhibitor), PIK-93 (PI3K inhibitor), Erlotinib (EGFR inhibitor), and Thalidomide (immunomodulatory drug). **2E.** The overlap between genes defining TNBC cell line identities and MsigDB hallmark gene sets. Bar plots show the percentage of overlapping genes across hallmark sets, with notable overlaps including EMT, TNFα signaling via NF-κB, estrogen responses (early and late), KRAS signaling, and interferon responses. These findings further validate the biological significance of the identified identities and their connections to known pathways. **2F.** The enriched drug-like molecule classes that show significant differential responses between TNBC and other cancer types. Together, these panels demonstrate the utility of integrating multi-omics data to define and validate biologically relevant subpopulation identities within TNBC cell lines.

We analyzed gene expression (GE) data to identify the most significantly differentially expressed genes in each subpopulation compared to other subpopulations. Some of the most significant genes were independently validated, confirming their biological relevance. For example, in the F3 subpopulation, genes associated with EGFR signaling were significantly upregulated (**Fig. 2B**) and MYC expression is associated with F1 identity (**Fig. 2B**).

We examined CRISPR loss-of-function screen data to identify unique gene dependencies of specific subpopulations. Notably, in the F2 and F3 subpopulations, PIK3CA and EGFR emerged as the top genes showing significant differential effects (**Fig. 2C**). These findings align with the biological identities of these subpopulations: F2 is associated with luminal-like and PI3K-Akt signaling signatures, while F3 is associated with EGFR and NF-κB signaling. In addition, the F0 subpopulation demonstrated dependencies on RBPJ and MIB1, genes critical to Notch signaling^49,50^, further reinforcing the link between this subpopulation’s identity and its molecular characteristics.

Using large-scale population-wide drug sensitivity curves from GDSC and CTRP, we investigated drugs whose efficacy varied according to the “weight” or explained variance score derived from the NMF matrix, which quantifies the association of each cell line with a given subpopulation identity. Because the F2 identity is characterized by PI3K-Akt signaling and PI3K gene dependency, we hypothesized that cell lines with increasing association with F2 identity would have enhanced sensitivity to the PI3K and Akt inhibitors. As predicted, we observed correlation between F2 identity and sensitivity to PI3K inhibitors, such as PIK-93 (**Fig. 2D and Extended Data** Fig.2A**,B**) and Akt inhibitors, Afuresertib and Uprosertib (**Extended Data** Fig.2C**,D**). Additionally, we observed correlations between F3 identity and sensitivity to EGFR inhibitors, namely erlotinib (**Fig. 2D and Extended Data** Fig.2E**,F**). These findings suggest that subpopulation identities derived from highly variable gene expression levels directly associate with drug response *in vitro*.

To further validate the biological context of these identities, we compared their defining genes to hallmark gene sets from MsigDB^51^. For some identities, 10–15% of genes also appear in key MsigDB hallmark gene sets (**Fig. 2E**). For example, the F2 identity strongly overlaps with hallmark gene sets related to estrogen response (early and late), consistent with its luminal-like characteristics (**Fig. 2E**). The F2 identity also overlaps with the androgen response gene set, which is consistent with the luminal androgen receptor subtype of TNBC^52–54^. Importantly, our framework offers insights that go beyond those provided by hallmark gene sets; hallmark gene sets are typically derived from single perturbation experiments, often using controlled cell line models subjected to specific interventions, such as CRISPR knockouts or drug treatments. These sets capture changes in gene expression or chromatin states driven by the dynamics of the experimental system. In contrast, our approach provides a global, population-wide view of heterogeneity, identifying gene “signatures” that reflect variability across the population rather than within the context of a specific experimental perturbation.

Finally, we analyzed drug response data in TNBC cell lines from CCLE, GDSC, and CTRP to identify classes of drugs with significantly different effects in TNBC compared to other cancer types. Out of 812 drugs, epigenetic remodeling agents stood out as the only class showing significant enrichment and efficacy in TNBC cell lines (**Fig. 2F**). These agents modify chromatin accessibility, influencing gene expression, and are increasingly being explored for their therapeutic potential in TNBC^55,56^. Notably, the F1 identity exhibited a significant sensitivity to CBB-1007, a histone demethylase inhibitor, further underscoring the relevance of epigenetic regulation in this subpopulation (**Fig. 2D**). The enrichment of epigenetic remodeling agents in TNBC highlights their promise for targeting this aggressive and therapeutically challenging cancer subtype.

### Deriving Single-Cell Identities and Tracking Drug-Responsiveness

To connect bulk-derived subpopulation identities to single-cell resolution, we used a reverse non-negative matrix factorization (NMF) approach^57^ (**Fig. 3A**). This method enables the mapping of subpopulation identities derived from bulk RNA-seq data to individual cells within single-cell RNA-seq (scRNA-seq) datasets. Conceptually, the bulk RNA-seq data provides a framework for defining overarching subpopulation identities, represented as a matrix of gene expression patterns (W matrix). For a given single-cell dataset, we input its gene expression matrix (GE matrix) and solve for the contribution of each subpopulation (H matrix). Here, the W matrix comes from bulk RNA-seq data from CCLE TNBC cell lines, and the H matrix represents the single-cell labels assigned to each subpopulation (**Fig. 3A**). This allows us to annotate each cell in a single-cell dataset with a specific identity from the bulk-derived framework (**Extended Data** Fig.3).

**Figure 3.**
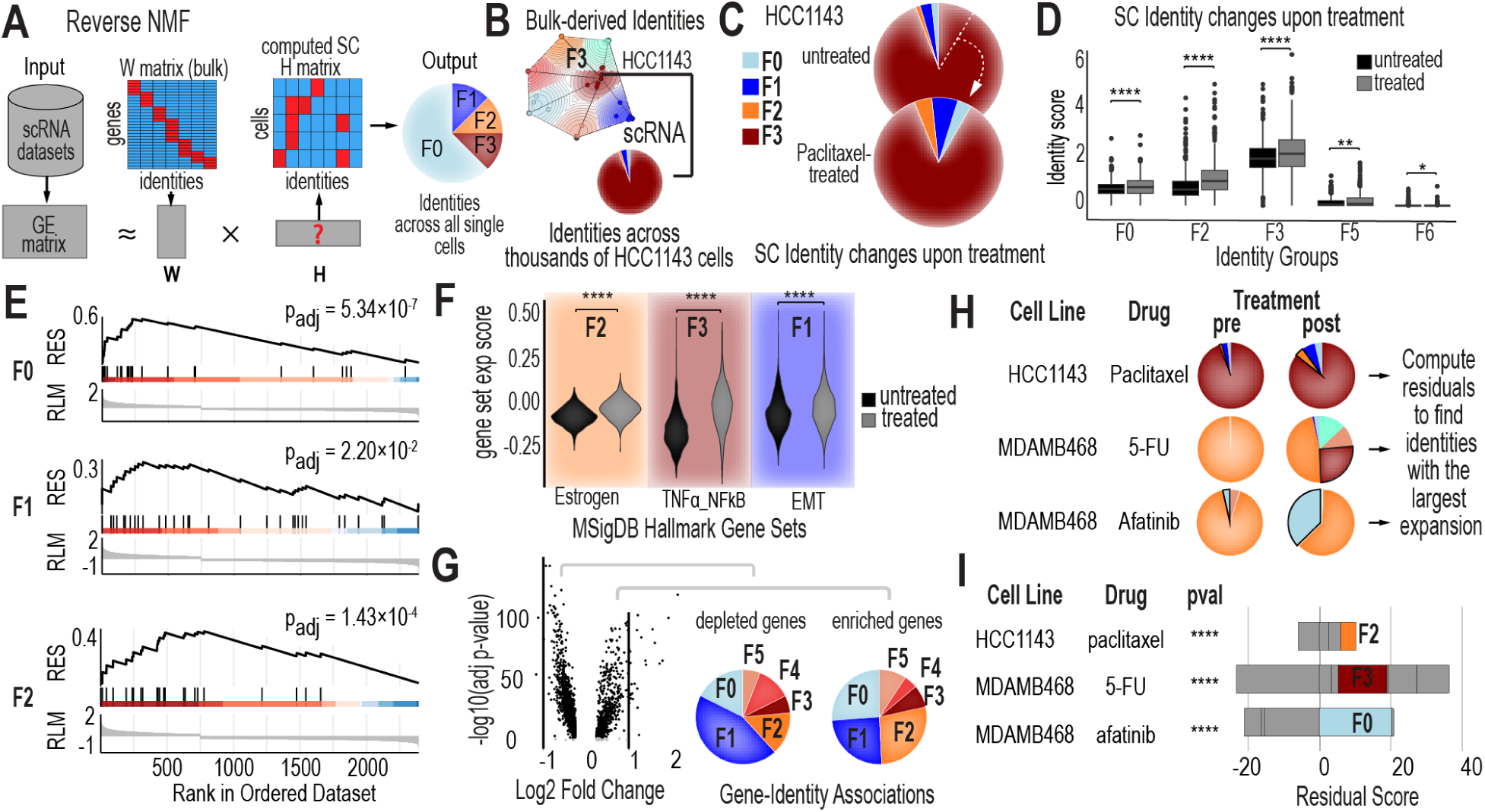
Characterizing Cellular Identity Changes in TNBC Cell Lines Upon Drug Treatment. **3A.** Workflow illustrating the mapping of bulk-level cancer cell line identities onto single cells in TNBC cell lines, enabling single-cell-level analysis of identity changes. **3B.** Single-cell identity composition of HCC1143, a TNBC cell line associated with the F3 identity in the CCLE model. The pie chart displays the distribution of single cells across different identities within this cell line prior to treatment. **3C.** Single-cell identity shifts in HCC1143 after paclitaxel treatment. Pie charts illustrate the percentage of single cells belonging to each identity before and after treatment, revealing changes in the identity composition. **3D.** Identity scores for single cells in HCC1143 before and after paclitaxel treatment. These scores, derived from the reverse NMF H matrix, indicate the intensity of gene expression identities at the single-cell level. **3E.** Gene Set Enrichment Analysis (GSEA) results for F0, F1, and F2 gene sets after paclitaxel treatment. The analysis highlights the enrichment or depletion of these gene sets in treated single cells, providing insights into identity-specific responses to treatment. **3F.** Expression scores for three MsigDB hallmark gene sets in HCC1143 single cells before and after treatment. The hallmark gene sets analyzed include estrogen response (early and late combined), TNFα signaling via NF-κB, and epithelial-mesenchymal transition (EMT), showcasing treatment-induced changes in key biological pathways. **3G.** Volcano plots of differentially expressed genes (DEGs) in HCC1143 single cells after paclitaxel treatment. For upregulated genes, the pie chart on the right illustrates the proportion of DEGs belonging to each identity as defined in the CCLE model. Similarly, for downregulated genes, the pie chart on the left shows the identity distribution of DEGs. **3H.** Illustration of residual score computation to quantify identity changes after drug treatments. Residual scores provide a measure of how much each identity expands or contracts in response to treatment. **3I.** Global characterization of identity changes across different TNBC cell lines treated with various drugs/compounds. Chi-squared test p-values assess the significance of identity changes for each cell line after treatment. Stacked bar plots depict the residual scores for each identity, with positive residual scores indicating identity expansion and negative residual scores indicating identity shrinkage.

As a proof-of-principle case study, we analyzed scRNA-seq data from paclitaxel-treated and untreated HCC1143 cells. Using our framework, we mapped individual cells to their closest matching identity, revealing that 94% of the HCC1143 population belonged to the F3 subpopulation, while 6% were distributed among F0, F1, and F2 identities (2% F0, 3% F1, 1% F2). The F3 subpopulation is characterized by distinct molecular features, including upregulated EGFR and NF-κB signaling (**Fig. 3B**). Notably, the predominance of F3 cells aligns with unmatched bulk RNA-seq data from the same cell line, where HCC1143 is classified as having an F3 identity at the bulk level due to the dominance of this subpopulation.

Paclitaxel treatment caused a notable expansion of the non-F3 subpopulations, increasing overall population heterogeneity (**Fig. 3C**). Subpopulation identity scores before and after treatment revealed significant shifts in most identities (**Fig. 3D**), demonstrating how drug treatment reshapes the population’s identity composition, with nearly all identity scores showing dynamic changes.

To validate the significance of these shifts at the gene level and understand their impact on the population as a whole, we performed gene set enrichment analysis (GSEA) on the scRNA-seq data. GSEA provided quantitative metrics, revealing significant enrichment and depletion of gene sets associated with distinct subpopulation identities. This analysis confirmed that genes associated with the F0, F1, and F2 identities were significantly enriched following drug treatment (**Fig. 3E**). Additionally, hallmark gene sets from MsigDB most closely aligned with these expanding subpopulations—such as estrogen response for F2, TNFα signaling via NF-κB for F3, and EMT for F1—exhibited significant changes, providing complementary evidence of the biological relevance of these identities and their treatment-induced shifts (**Fig. 3F**). These findings highlight how these subpopulations are linked to distinct biological processes that are dynamically regulated in response to treatment.

We aimed to determine whether specific genes disproportionately contributed to identity shifts during drug treatment. To address this, we performed differential gene expression analysis between treated and untreated conditions (**Fig. 3G**), identifying the most significantly altered genes and mapping them to their associated subpopulation identities. While gene-level changes were observed across all identities, the F0, F2, and F3 subpopulations exhibited the most substantial changes in HCC1143, suggesting a potential role for these genes and identities in drug responsiveness. Our analysis further revealed that certain identities played a dominant role in driving the observed shifts. For instance, in HCC1143, the F2 identity emerged as the primary driver of change, contributing most significantly to the overall subpopulation-level shifts observed after treatment (**Fig. 3I**). Extending this analysis to other TNBC cell lines (**Fig. 3H**) confirmed that specific identities, such as F2 and F3, were consistently associated with the most adaptive and dynamic subpopulations (**Fig. 3I and Extended Data Fig.4**).

Our proof-of-principle analysis demonstrates the strength of integrating bulk-derived subpopulation identities with single-cell data to uncover how distinct subpopulations shift and drive adaptation during drug treatment. By defining cell populations through subpopulation-level identities, we show that certain identities—or their combinations—are closely linked to drug responsiveness, offering valuable insights into the population-level mechanisms underlying treatment adaptation.

### Decoding Drug Response in Patient Samples Using Single-Cell Identity Annotation

We applied our framework to single-cell transcriptomics data from 13 TNBC patient samples treated with the anti-PD1 immune checkpoint inhibitor pembrolizumab^58^. Of these, 11 were treatment-naive prior to pembrolizumab, while 2 (BIOKEY_41, B41, and BIOKEY_35, B35) received neoadjuvant chemotherapy before anti-PD1 treatment. Single-cell analyses grouped cells into distinct clusters based on gene expression, corresponding to distinct cell types, with drug treatment inducing noticeable shifts within these clusters (**Fig. 4A**). Using our bulk-derived identity mapping framework, we mapped each cancer cell to its corresponding identity across all treatment-naive and treated patient samples. This enabled us to characterize the composition of tumor subpopulations within each sample and track how these compositions changed after treatment. By comparing pre- and post-treatment compositions, we identified subpopulations with unique identity profiles that were either enriched, depleted, or exhibited adaptive responses to pembrolizumab.

**Figure 4.**
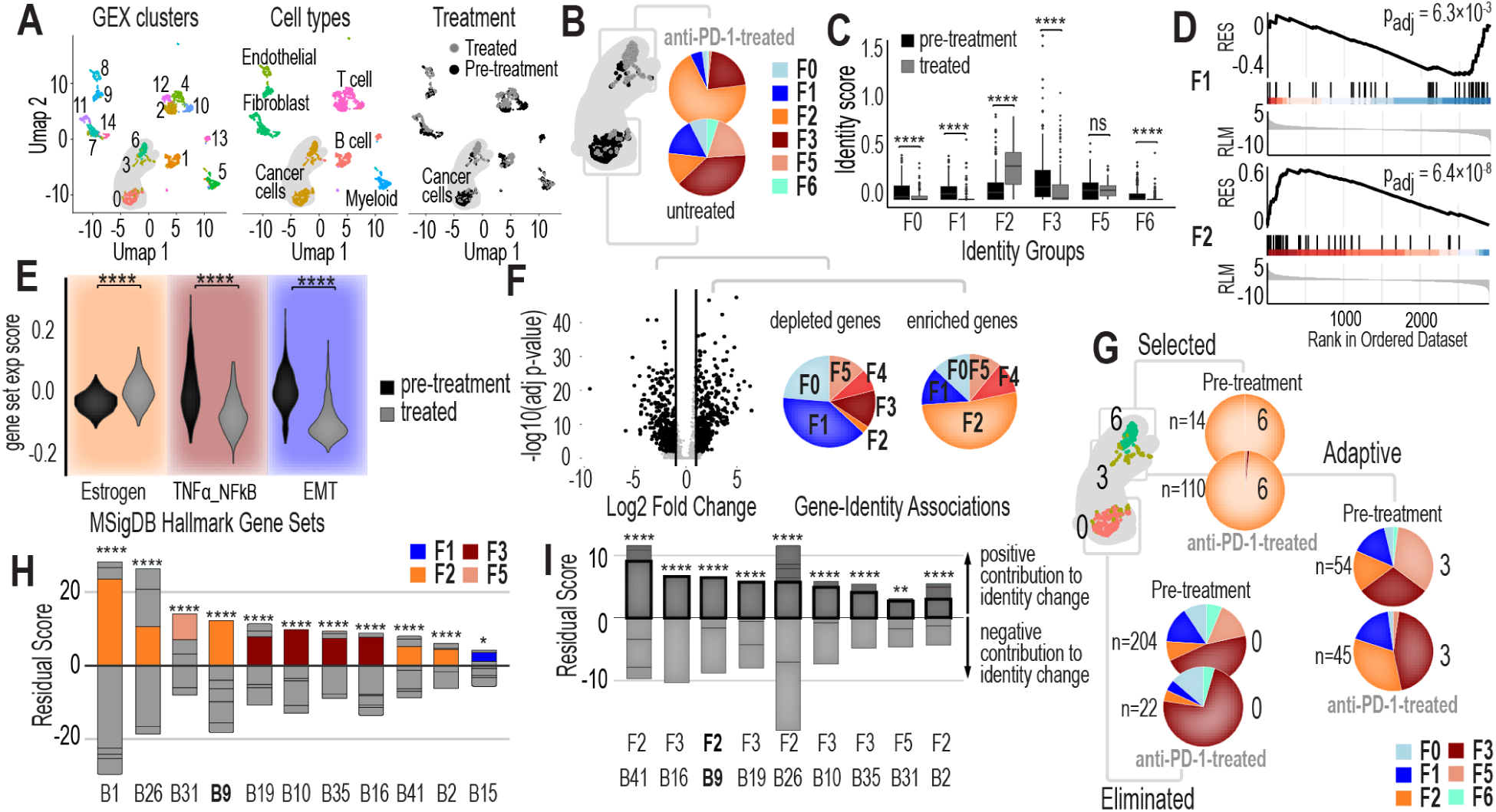
Characterizing Cellular Identity Changes in TNBC Tumor Cancer Cells Upon Pembrolizumab Treatment Using the CCLE Tumor-Intrinsic Model. **4A.** UMAP plots showing cluster identifications, cell types, and treatment conditions for all single cells in the BIOKEY_9 (B9) tumor. These plots provide an overview of the cellular landscape within the tumor, highlighting distinct clusters and their associations with treatment conditions. **4B.** Single-cell identity changes in the cancer cells of TNBC patient BIOKEY_9 (B9) after pembrolizumab treatment. Pie charts illustrate the distribution of single cells across different identities within the cancer cell population before and after treatment. **4C.** Identity scores for single cells in the BIOKEY_9 (B9) cancer cell population before and after pembrolizumab treatment. These scores quantify the intensity of gene expression identities at the single-cell level, revealing shifts in cellular identities induced by treatment. **4D.** GSEA results for F1 and F2 gene sets after pembrolizumab treatment, highlighting the enrichment or depletion of these identity-specific gene sets in the treated cancer cell population. **4E.** Expression scores for three MsigDB hallmark gene sets in BIOKEY_9 (B9) cancer single cells before and after treatment. The hallmark gene sets analyzed include estrogen response (early and late combined), TNFα signaling via NF-κB, and epithelial-mesenchymal transition (EMT), providing insights into treatment-induced changes in key biological pathways. **4F.** Volcano plots of differentially expressed genes (DEGs) in BIOKEY_9 (B9) cancer single cells after pembrolizumab treatment. Pie charts accompanying the plots display the percentage of DEGs associated with each identity in the CCLE model. The right pie chart represents upregulated genes, while the left represents downregulated genes, linking these changes to specific identities. **4G.** Analysis of identity composition changes in three cancer cell clusters from BIOKEY_9, B9, (cluster_0, cluster_6, and cluster_3) before and after pembrolizumab treatment. Pie charts illustrate the percentage of single cells belonging to each identity within each cluster, revealing differential responses across clusters. **4H.** Global characterization of identity enrichment or depletion for cancer cells within 11 TNBC tumors following anti-PD1 treatment. Chi-squared test results assess the significance of identity changes across tumors, with stacked bar plots depicting residual scores for each identity. Positive residual scores indicate identity expansion, while negative scores indicate depletion. **4I.** Analysis of cancer cell clusters contributing to the largest identity expansions in cancer cells for each tumor. Chi-squared test p-values assess whether specific clusters are more associated with identity expansion compared to other clusters within the same tumor. Stacked bar plots represent the residual scores for each cluster, highlighting their contributions to identity changes.

As a proof-of-concept, we applied our approach to a representative patient sample (BIOKEY_9) with tumor samples collected before and after pembrolizumab monotherapy (**Fig. 4A**). We identified six distinct subpopulations within the tumor cells, each composed of different identities (**Fig. 4B**). Prior to treatment, the majority of cells were part of the F3 subpopulation; however, following treatment, the F2 subpopulation became predominant. The most notable shifts involved the expansion of F2 and the depletion of F1 and F3 identities (**Fig. 4C**). Gene set enrichment analysis^59^ (GSEA) further validated the significance of the shifts in F1 and F2 identities (**Fig. 4D**).

These identity shifts correspond to biologically relevant changes in MsigDB hallmark gene sets associated with each identity. For example, the F2 identity exhibited a significant increase in estrogen response (early and late), the F3 identity showed a decrease in TNFα signaling via NF-κB, and the F1 identity displayed depletion of EMT-related processes (**Fig. 4E**). These findings align with the known biological roles of these subpopulations: F2 is linked to luminal-like characteristics, while F3 is associated with TNFα signaling via NF-κB and EGFR pathways. Differential gene expression analysis further revealed that most genes enriched after treatment mapped to the F2 identity, while the majority of depleted genes were associated with the F1 identity (**Fig. 4F**).

We sought to determine whether distinct clusters of cells, initially grouped by gene expression, exhibited unique identity shifts in response to drug treatment. By analyzing these clusters independently, we uncovered striking differences in their behavior following treatment (**Fig. 4G**). In the first cluster (cluster_6), predominantly composed of cells with an F2 identity, the identity composition remained stable, but the total number of cells increased after treatment, suggesting a selection mechanism favoring this subpopulation. In contrast, the second cluster (cluster_0), a mix of six identities dominated by F3, experienced a dramatic decline in cell numbers post-treatment, indicating possible elimination or selection against this subpopulation. The third cluster (cluster_3), another mix of six identities, displayed substantial shifts in composition without a significant change in total cell numbers, pointing to an adaptive subpopulation capable of dynamically reconfiguring under drug pressure. These findings highlight the power of our framework in dissecting drug responsiveness, revealing how distinct cell clusters, defined by their unique identity compositions, contribute differently to treatment outcomes.

### Global Patterns of Adaptation Across Patient Samples

Building on these findings, we expanded our analysis to multiple patient samples^58^ (**Extended Data Fig.5,6 and Supplementary Table 7**) to determine whether global trends in identity shifts could be identified. This analysis revealed an intriguing pattern: certain identities consistently played a disproportionate role in driving changes in subpopulation composition during treatment (**Fig. 4H**). To quantify these contributions, we calculated a residual score for each identity, measuring the extent to which they influenced the overall shifts observed (**Fig. 4H**). To investigate if disproportionate contribution to identity shifts from certain cancer cell clusters is a global trend, we focused on the identity with the most significant expansion during treatment for each patient and calculated a residual score for each cancer cell cluster regarding the identity expansion (**Fig. 4I**). This approach enabled us to capture both positive and negative contributions from each cluster to the identity expansion, with identities with the most significant expansion during treatment highlighted in the figure (**Fig. 4I**). Globally, we found that distinct cell clusters seemed to contribute differently to molecular treatment outcomes.

Notably, the F2 and F3 identities consistently emerged as key drivers of the largest shifts in subpopulation composition across patient samples (**Fig. 4H**). Subpopulations, grouped by gene expression and defined by their identity composition, revealed that those dominated by F2 and F3 identities exhibited the most pronounced changes in response to drug treatment. These shifts, characterized by significant expansion or decline of these identities, highlight their pivotal role in driving treatment-induced adaptation. These findings emphasize that the composition of subpopulations—particularly their enrichment in F2 and F3 identities—is a critical determinant of therapy responsiveness, underscoring the value of identity-level analysis in understanding treatment outcomes.

These findings underscore the importance of identifying and targeting adaptive subpopulations at the single-cell level. Building on this, future studies may explore whether tumors enriched in F2 and F3 identities possess an enhanced capacity to adapt and evade therapy, driven by their distinct biological characteristics. The F2 identity is linked to luminal-like traits and PI3K-Akt signaling, while F3 is associated with NF-κB and EGFR pathways—hallmarks of treatment resistance and adaptability. The disproportionate influence of these identities highlights how tumors leverage specific adaptive programs under therapeutic pressure. By targeting these dominant subpopulations, it may be possible to disrupt critical survival and adaptive mechanisms, offering a promising strategy to mitigate therapy resistance and improve long-term treatment outcomes.

### Modeling Complexity: Identifying Drug-Responsive Identities in a TNBC Organoid Model

Our findings reveal that certain identities are inherently more adaptive, driving significant shifts that shape therapeutic adaptation. To investigate the mechanisms underlying these shifts, we developed an organoid model system that captures the complexity of tumor heterogeneity while providing a testable framework for assessing drug sensitivity. Unlike traditional cell lines, organoid models offer a more realistic representation of tumor diversity, enabling us to integrate single-cell RNA sequencing with single-cell epigenetic profiling to study treatment responses. Applying our framework to these organoid models allowed us to validate the identity shifts observed during treatment and uncover the molecular and epigenetic mechanisms driving these adaptations.

We developed an organoid model derived from a primary human TNBC tumor obtained from a UNC Lineberger Comprehensive Cancer Center patient (**Extended Data Fig.7**) and treated it with the epigenetic remodeling agents JQ1 (a BRD4 inhibitor) and MS177 (an EZH2 degrader). These agents were selected based on prior findings (**Fig. 2F**) showing that epigenetic remodeling drugs elicited the most significant differences in drug response between TNBC cell lines and other cancer types. The organoid model demonstrated sensitivity to both drugs (**Fig. 5F and Extended Data Fig.9A**). To further investigate these responses, we generated matched single-cell multiomics data before and after treatment with each drug and applied our multimodal framework to analyze subpopulation identity shifts and their role in treatment adaptation.

**Figure 5.**
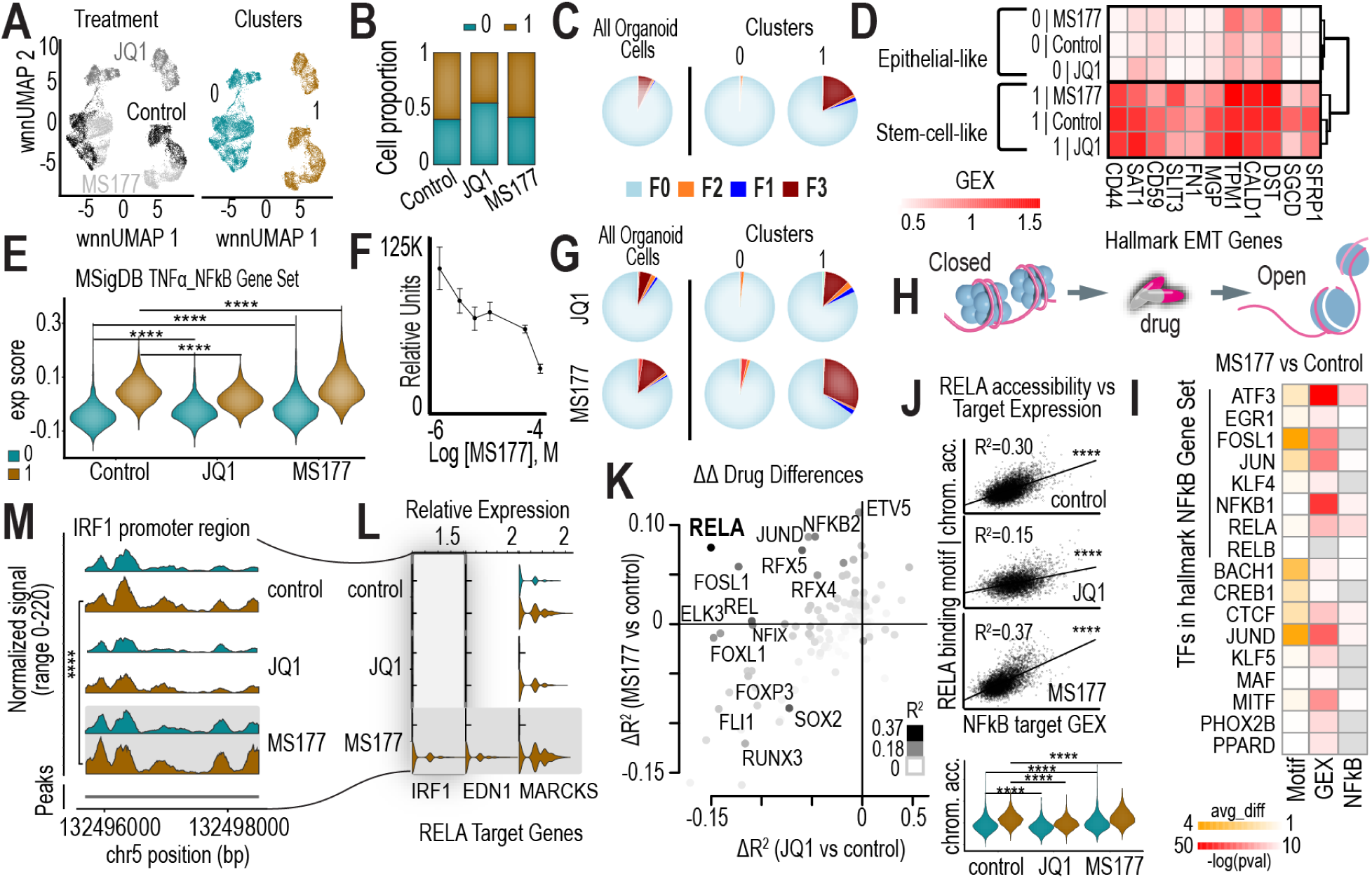
Investigating Identity Changes After Drug Treatment in a TNBC Organoid Model. **5A.** UMAP plots showing treatment conditions and cluster identifications of single cells within the TNBC organoid, providing an overview of the cellular landscape under different treatment conditions. **5B.** Bar plots depicting the proportions of single cells belonging to each cluster across the three treatment conditions, highlighting shifts in cluster composition following drug treatment. **5C.** Pie charts illustrating the percentage of single cells belonging to each identity in the organoid as a whole and within each of the two distinct clusters under control conditions. **5D.** Comparison of gene expression levels for stem-cell/EMT markers between the epithelial-like and stem-cell-like clusters within the organoid across the three treatment conditions, revealing identity-specific transcriptional profiles. **5E.** Expression scores for the MsigDB hallmark gene set “TNFα signaling via NF-κB” for the two clusters within the organoid across the three treatment conditions, demonstrating differential pathway activation. **5F.** Drug treatment curve showing the response of the organoid to MS177, indicating the relative efficacy of the treatment on the organoid population. **5G.** Pie charts showing the percentage of single cells belonging to each identity in the organoid as a whole and within the two distinct clusters under JQ1 and MS177 treatment conditions. **5H.** Schematic illustration depicting how drug treatments, such as MS177, open chromatin regions, enabling potential gene transcription and altering regulatory dynamics. **5I.** Analysis of transcription factors (TFs) relevant to the response to MS177 treatment. The first column shows the average difference in TF binding motif accessibility after MS177 treatment. The second column highlights the significance of TFs in upregulating gene expression broadly after MS177 treatment. The third column focuses on TFs that significantly upregulate gene expression specifically in NF-κB pathway genes after MS177 treatment. **5J.** Scatterplots showing R^2^ between RELA binding motif accessibility and the expression scores of RELA target genes in the NF-κB pathway under control, JQ1-treated, and MS177-treated conditions. Violin plots below display RELA binding motif chromatin accessibility levels in the two clusters (clusters 0 and 1) across the three treatment conditions. **5K.** Scatterplot comparing changes in R² values between TF binding motif accessibility and the expression scores of their target genes in the NF-κB pathway following JQ1 (x-axis) and MS177 (y-axis) treatments for all TFs. Dot colors indicate the R² value under MS177-treated conditions. **5L.** Gene expression levels of RELA targets MARCKS, EDN1, and IRF1 in the two clusters across the three treatment conditions. Chromatin accessibility in the promoter regions of these genes, where RELA binds, increases after MS177 treatment, linking chromatin remodeling to transcriptional changes. **5M.** Chromatin accessibility of the IRF1 promoter region in the two clusters across the three treatment conditions, demonstrating treatment-induced epigenetic remodeling.

Both scRNA and scATAC sequencing analyses revealed distinct drug-treatment clusters and corresponding gene expression and accessibility clusters, labeled 0 and 1 (**Fig. 5A**). Across all three treatment conditions, these clusters maintained a consistent 3:2 cell number ratio (**Fig. 5B**). Mapping cells to bulk-derived CCLE subpopulation identities showed that the organoid was predominantly represented by the F0 state, characterized by partial EMT and Notch signaling. Notably, the identity compositions of the two clusters differed significantly: Cluster 0 was almost entirely dominated by the F0 identity, while nearly one-quarter of the cells in Cluster 1 belonged to F1, F2, and F3 subpopulations, highlighting a greater degree of identity heterogeneity (**Fig. 5C**).

Given the presence of the F1 identity in cluster 1, we analyzed hallmark EMT gene sets under control and treatment conditions, revealing that cluster 1 exhibited a stem-cell-like phenotype, while cluster 0 was more epithelial-like (**Fig. 5D**). Similarly, the F3 identity in cluster 1 prompted an analysis of hallmark TNFα via NF-κB signaling gene sets, which showed significant differences between the epithelial (low NF-κB) and stem-like (high NF-κB) clusters. Notably, MS177 treatment specifically increased NF-κB signaling in cluster 1 cells, further reinforcing its stem-like characteristics^60,61^ (**Fig. 5E**). Remarkably, our analysis uncovered the adaptive nature of cluster 1 identities during drug treatment. For example, the F3 identity expanded to nearly one-third of the total cell population following MS177 treatment, emphasizing its adaptive capacity (**Fig. 5G**).

### Drug-Responsive Identities Are Defined by Distinct Epigenetic Regulation

We hypothesized that the large-scale identity shifts observed in cluster 1 during drug treatment are driven by significant changes in chromatin accessibility. Chromatin accessibility, which determines how “open” or “closed” DNA is at specific loci, directly regulates the binding of transcription factors and other regulatory proteins, thereby influencing gene expression (**Fig. 5H**). With epigenetic remodeling drugs, we anticipate substantial changes in chromatin accessibility that ultimately drive the identity composition shifts within this cluster. For instance, in cluster 1 cells, altered chromatin accessibility at key regulatory regions likely corresponds to the gene expression shifts observed in identities such as F1, F2, and F3, which significantly expand following MS177 treatment.

To test this hypothesis, we analyzed chromatin accessibility changes at transcription factor (TF) binding motifs associated with regulators of the NF-κB pathway. Specifically, we examined TF binding motif accessibility changes after MS177 treatment (**Fig. 5I**, first column), assessed the general involvement of these TFs in upregulating gene expression (**Fig. 5I**, second column), and evaluated their specific roles in the NF-κB signaling pathway (**Fig. 5I**, third column). Several transcription factors, including ATF3 and RELA, exhibited increased binding motif accessibility alongside upregulated gene expression in the NF-κB signaling pathway, implicating these TFs as key mediators of MS177-induced transcriptional changes. Notably, RELA, a critical NF-κB subunit required for nuclear translocation and activation, emerged as a compelling candidate, with its binding motif accessibility strongly correlating with the expression of its target genes in the NF-κB signaling pathway (**Fig. 5J**).

We further examined the relationship between chromatin accessibility and gene expression by systematically analyzing the correlation between accessibility at enriched transcription factor binding motifs and the expression of the transcription factor’s target genes within the NF-κB pathway. Consistent with expectations, RELA demonstrated the strongest correlation (R²) in the MS177-treated condition (**Fig. 5J,K, Extended Data Fig.8, and Extended Data Fig.9B-D**), with this relationship significantly enhanced under MS177 treatment (**Fig. 5J,K**). Furthermore, RELA binding motif accessibility was specifically induced by MS177, with a higher level observed in cluster 1 cells (**Fig. 5J**, bottom panel). These findings establish a mechanistic link between RELA activity, the expansion of NF-κB signaling, and the adaptive responses exhibited by cluster 1 cells during MS177 treatment.

To investigate how these changes correlate with specific subpopulation identities, we focused on transcription factors with selective increases in R² under MS177 treatment but not JQ1. RELA again emerged as a key regulator (**Fig. 5K**), aligning with the observed expansion of the F3 identity, which occurs predominantly under MS177 treatment. Within this framework, three RELA target genes in the NF-κB pathway—IRF1, EDN1, and MARCKS—exhibited distinct increases in gene expression, consistent with the F3 identity expansion under MS177 treatment (**Fig. 5L**). These genes, along with other genes in the NF-κB hallmark pathway (**Extended Data Fig.10**), displayed higher expression in cluster 1 compared to cluster 0, specifically in response to MS177. Additionally, chromatin accessibility at the promoter regions of these three genes increased following MS177 treatment (**Supplementary Table 8–10**). These findings further support the role of RELA in mediating NF-κB signaling and the adaptive expansion of the F3 subpopulation during treatment.

We investigated the promoter regions of these genes for chromatin accessibility differences. For IRF1, we observed significant accessibility differences between cluster 0 and cluster 1, particularly after MS177 treatment (**Fig. 5M**). This observation suggests that epigenetic regulation plays a significant role in shaping the F3 identity during MS177 treatment. While IRF1 promoter accessibility provides an illustrative example of these changes, it likely reflects broader epigenetic differences between clusters that contribute to the dynamic remodeling of subpopulation identities during drug treatment.

Our analyses reveal that the adaptive changes in cluster 1 may be explained by increased NF-κB signaling and chromatin remodeling, which collectively support the expansion of the F3 identity. These findings highlight that drug-responsive identities, such as F3, are likely shaped not only by transcriptional changes but also by dynamic shifts in epigenetic regulation. The observed differences in chromatin accessibility between clusters, alongside transcription factor activity, uncover key adaptive mechanisms driving subpopulation shifts under therapeutic pressure. This work underscores the critical interplay between epigenetic and transcriptional regulation in defining drug-responsive identities and emphasizes the potential of targeting these mechanisms to disrupt tumor adaptation. By linking changes in gene expression with chromatin accessibility, we gain valuable insights into how cancer subpopulations adapt, paving the way for more effective therapies that exploit the vulnerabilities of cells driven by adaptive identities.

## Conclusion

This study introduces a novel multimodal framework that integrates bulk and single-cell transcriptomics to define and characterize drug-responsive subpopulation identities. By bridging large-scale population data with single-cell resolution, this approach uncovers meaningful biological signals within the complexity of single-cell data. It enables systematic mapping of adaptive cell identities under therapeutic pressure, addressing a key challenge in cancer research and advancing the development of precise strategies to overcome treatment resistance.

Through our analyses of cell lines, organoid models, and patient samples, we discovered global patterns of drug response that consistently point to the adaptive response of select subpopulation identities. Across these systems, certain identities emerge as more malleable, driving significant shifts in subpopulation identity composition during drug treatment. These findings emphasize the importance of understanding how tumors evolve at the level of their cellular diversity, providing key insights into the mechanisms that enable therapeutic evasion.

Our work also highlights the utility of single-cell multiomics in explaining the observed shifts in gene expression with changes in epigenetic regulation. The ability to link these shifts underscores the utility of single-cell multiomics in revealing the molecular underpinnings of adaptive subpopulations. Here, we discover broader epigenetic differences exist between clusters and contribute to the dynamic remodeling of subpopulation identities during epigenetic remodeling drug treatment.

This framework contributes to the growing body of work that seeks to unravel the complexities of cancer heterogeneity and treatment response. By enabling the identification of adaptive subpopulations and their underlying mechanisms, it provides a critical step toward precision medicine. Ultimately, this work lays the foundation for therapeutic strategies that not only target tumors at their current state but also anticipate and disrupt their future adaptive trajectories, offering new hope for more durable and effective cancer treatments.

## Data Availability

All scripts are available at GitHub (https://github.com/Brunk-Lab/TNBC_Identities). Single cell multiomics data is available on Gene Expression Omnibus (GEO accession number GSE288798).

## Methods

### Data Sources and Processing

**TNBC Cell Line Data:**

- **RNA-Seq Data:** Processed RNA sequencing data (v22Q2) for Triple Negative Breast Cancer (TNBC) cell lines were obtained from the Cancer Cell Line Encyclopedia (CCLE) via the Dependency Map (DepMap) portal.
- **CRISPR Gene Effect Data:** CRISPR gene dependency scores (v22Q2) were sourced from CCLE.
- **Drug Sensitivity Data:** Drug sensitivity datasets from the Cancer Therapeutics Response Portal (CTRP) (v2.0) and the Genomics of Drug Sensitivity in Cancer (GDSC) were also accessed via the CCLE/DepMap portal.

**Patient Tumor Data:**

- **TCGA (**The Cancer Genome Atlas**) Data:** Processed RNA sequencing data and survival data for breast cancer tumors were retrieved from the study Breast Invasive Carcinoma (TCGA, PanCancer Atlas) (brca_tcga_pan_can_atlas_2018) from cBioPortal. For basal-like breast cancer (BLBC) analysis, PAM50 intrinsic subtypes were assigned to each patient sample using an ER/HER2 subgroup-specific gene-centering method^62^. Basal subtype samples were selected for further analysis.

**Single-Cell RNA-Seq Data:**

- **TNBC Cell Lines (Treatment-Naive):** scRNA-seq data for treatment-naive TNBC cell lines were retrieved from GEO datasets GSE182694, GSE174391, GSE152315, GSE173634, and GSE202771.
- **TNBC Cell Lines (Drug-Treated):** scRNA-seq data for TNBC cell lines under drug treatment were sourced from GEO datasets GSE131135, GSE139129, GSE164716, and GSE228154.

- **HCC1143 (GSE139129):** Treated with paclitaxel for 24 and 72 hours.
- **MDAMB468 (GSE164715):** Treated with 5-fluorouracil (5-FU) for 33, 50, 67, 77, 171, 202, and 214 days.
- **MDAMB468 (GSE228154):** Treated with Afatinib for 3, 6, and 9 days.
- **TNBC Patient Tumors (Treatment-Naive):** scRNA-seq data accessed from GEO dataset GSE176078^63^.
- **TNBC Patient Tumors (Anti-PD1 Treatment):** scRNA-seq data from TNBC patients receiving anti-PD1 therapy were obtained from the Lambrechts Lab resource (http://biokey.lambrechtslab.org).

### Defining Gene Expression Landscapes and Identity Profiles in TNBC Cell Lines and Tumors

#### Gene Selection with High Variance Across Samples

To identify genes with high expression variance, we analyzed TNBC cell lines (CCLE) and basal-like breast cancer patient tumors (TCGA) separately. To further explore phenotypes associated with the MYC oncogene, DBSCAN clustering (Python sklearn package v1.2.2) was applied to two-dimensional expression data consisting of each gene and MYC. The eps parameter was systematically varied from 0.1 to 1.21 at intervals of 0.01 to optimize clustering. Genes were filtered based on DBSCAN^64^ cluster results.

For CCLE cell lines, genes with maximum expression values among TNBC cell lines lower than 3 were filtered out. Genes with expression ranges among TNBC cell lines lower than 3 were filtered out. Genes were selected if there was at least one cluster pair containing more than 3 TNBC cell lines in each cluster with cluster centroid distance above 3 and cluster separation above 1.

For TCGA patient samples, a first round of DBSCAN^64^ was run using all breast cancer samples. Then genes were selected for further filtering if cluster pair distances were more than 4 (higher than 95% of the quantile) and there were no less than 10 basal-like breast cancer tumor samples in each cluster of the pair. 753 genes were selected after the first-round filtering.

These 753 genes were sent for another round of DBSCAN running. In this round, only basal-like breast cancer tumor samples were used. Then genes were selected with cluster pairs that had centroid distance more than 4, and there were at least 8 basal-like breast cancer tumor samples in each cluster of the pair.

This process yielded 524 salient genes (including MYC) for TNBC cell lines and 489 salient genes (including MYC) for basal-like breast tumors.

#### Non-Negative Matrix Factorization (NMF) and Identity Mapping

The selected genes were used to perform NMF (Python ccal^65^ package v0.9.4) on both TNBC cell lines and tumor samples (with k=4–9). The elbow method determined the optimal k, after which hierarchical clustering (HC) was applied to the H and W matrices to group TNBC samples and genes, respectively. HC on the H matrix grouped TNBC samples into transcriptionally distinct identities, while the W matrix provided gene weights for each identity.

To annotate the biological relevance of each identity, top-expressed genes were identified, and an in-depth literature review was conducted (**Supplementary Table 1-6 and 11-15**). Common pathways or gene expression programs explaining these genes were assigned to define the biological characteristics of each identity. Using the ccal^65^ package, transcriptomic identity maps were generated for TNBC cell lines and basal-like breast tumor samples.

#### Multi-Omics Evidence Supporting Identity Biology

To validate the biological relevance of each identity, MsigDB hallmark gene sets were analyzed to identify genes supporting specific TNBC cell line subpopulations. For both hallmark and selected genes, expression levels between subpopulations and other samples were compared using Wilcoxon rank-sum tests, with adjusted p-values.

Essential genes for each subpopulation were identified using the CCLE CRISPR gene effect dataset. Gene effect scores were compared using Wilcoxon rank-sum tests.

#### Kaplan-Meier Survival Analysis

Kaplan-Meier survival curves were constructed for each identity of TCGA basal-like breast cancer patients using the KaplanMeierFitter() function from the Python lifelines package^66^ (v0.27.4). Survival comparisons between identities were performed using the Python logrank_test() function, with the number of subjects at risk displayed using add_at_risk_counts() from lifelines.plotting.

#### Linking High-Variance Genes and Subpopulation Identities to Drug Response

To evaluate whether the selected high-variance genes effectively capture molecular drug responses, we collected publicly available RNA-seq and scRNA-seq datasets from TNBC cell lines and tumors under various drug treatments (**Supplementary Table 16**). Differential gene expression (DGE) analysis was performed on these datasets. For scRNA-seq datasets, we focused on cancer cells to ensure tumor-specific insights. Enrichment of the selected high-variance genes within the differentially expressed genes (DEGs) was assessed using chi-squared tests, with results provided in **Supplementary Table 16**.

Additionally, we explored relationships between subpopulation identities and drug sensitivity by leveraging drug concentration-viability area under the curve (AUC) data from CTRP and GDSC. Pearson correlations were calculated between AUC values and the H matrix z-scores representing subpopulation-specific contributions. These analyses, conducted using pearsonr() from the scipy package^67^ (v1.8.0), highlight the degree to which certain identities influence drug response. Linear regression lines were computed with stats.linregress(). For drugs/compounds in these databases, AUC levels between subpopulations and other samples were compared using Wilcoxon rank-sum tests.

### Establishing and Treating a TNBC Organoid Model for Multiomics Analysis

Tumor samples were obtained from a triple-negative breast cancer (TNBC) patient at the University of North Carolina (UNC) under IRB-approved protocols. To ensure cell viability, the tumor tissue was processed within 1–2 hours of surgical resection. The tissue was finely minced and enzymatically digested using a collagenase-hyaluronidase mix (StemCell Technologies) for 1–2 hours at 37°C. Following digestion, red blood cells were removed using ammonium chloride solution, and the remaining cell clusters were resuspended in growth factor-reduced Matrigel (Corning). Cell clusters were seeded as 20 µL Matrigel domes in 6-well plates and cultured in breast organoid medium. This medium consisted of Advanced DMEM:F12 (Gibco) supplemented with B27, N2, Glutamax, HEPES, Penicillin/Streptomycin, Primocin, Nicotinamide, N-Acetylcysteine, R-spondin 3, Heregulin β-1, Noggin, FGF-10, FGF-7, EGF, A83-01, SB202190, and Y-27632 (added for the first three days to enhance cell survival). Media were refreshed every 3–4 days. Upon reaching confluency, typically once per week, organoids were manually dissociated and passaged by enzymatic digestion with TrypLE (Gibco) for 8 minutes at 37°C.

For drug treatment experiments, organoids were dissociated into single cells, counted, and 10,000 cells were resuspended in 10 µL of 70% Matrigel. These were seeded into 96-well plates as domes, solidified for 30 minutes, and overlaid with 100 µL of complete media. Organoids were allowed to form over 78 hours before drug treatment. Organoids were treated with various concentrations of JQ1 (a BRD4 inhibitor) and MS177 (an EZH2 degrader) for 48 hours, with DMSO serving as the diluent control, in triplicate. Viability was assessed using the CellTiter-Glo 3D Cell Viability Assay (Promega), which measures ATP levels as an indicator of viable cells. Luminescence was recorded using an Asteran microplate reader. For single-dose experiments, JQ1 and MS177 were administered at final concentrations of 500 nM and 250 nM, respectively, for 24 hours, after which organoids were harvested for downstream analyses.

Post-treatment, organoids were dissociated into single-cell suspensions using Accutase (Sigma) and gentle pipetting. Cells were filtered through a 40-µm strainer, washed with PBS, and counted with a Countess II Automated Cell Counter (ThermoFisher) to ensure >85% viability. Approximately 10,000 viable cells per condition were processed using the Chromium Single Cell Multiome ATAC + Gene Expression platform (10X Genomics). GEM generation, cDNA synthesis, and ATAC library preparation were conducted according to the manufacturer’s protocols, enabling downstream multiomic analysis of transcriptomic and chromatin accessibility changes.

### Organoid 10X Multiome Library Generation and Sequencing

To characterize the transcriptional and chromatin accessibility landscapes of TNBC organoids treated with JQ1 and MS177, we performed multiomics single-cell sequencing using the Chromium Single Cell Multiome ATAC + Gene Expression platform (10X Genomics). Organoids were harvested after 24 hours of drug treatment to capture cellular state changes while minimizing artifacts associated with extended culture. Single-cell suspensions were prepared following the 10X Genomics protocol. The cell concentration and quality were assessed using a Countess II Automated Cell Counter (ThermoFisher). Approximately 10,000 cells per sample were loaded into the Chromium Controller to partition individual cells into Gel Bead-In Emulsions (GEMs), enabling simultaneous profiling of RNA transcripts and chromatin accessibility within the same cells. Library preparation followed the manufacturer’s guidelines, including reverse transcription for gene expression, transposition for chromatin accessibility, and amplification of both cDNA and ATAC libraries. Libraries were quantified using a Qubit dsDNA High Sensitivity Assay (ThermoFisher) and assessed for fragment size distribution using an Agilent 4200 TapeStation. Sequencing was performed using the NextSeq2000 (Illumina) with paired-end reads to ensure high-resolution data.

### Multiomics Data Analysis

We conducted comprehensive analyses integrating scRNA-seq and single-cell multiome data to investigate transcriptional and chromatin accessibility changes in TNBC cell lines, organoids, and patient samples.

#### scRNA-seq Analysis

For scRNA-seq data from TNBC cell lines under drug treatment, we focused on HCC1143 treated with paclitaxel for 24 hours, MDAMB468 treated with 5-FU for 50 days, and MDAMB468 treated with Afatinib for 3 days. For scRNA-seq datasets of TNBC patients treated with anti-PD1 therapy, we included samples with sufficient cancer cell counts (IDs: BIOKEY_1 (B1), BIOKEY_2 (B2), BIOKEY_9 (B9), BIOKEY_10 (B10), BIOKEY_11 (B11), BIOKEY_14 (B14), BIOKEY_15 (B15), BIOKEY_16 (B16), BIOKEY_19 (B19), BIOKEY_26 (B26), BIOKEY_31 (B31), BIOKEY_35 (B35), BIOKEY_41 (B41)). For patient scRNA-seq data, cancer cells were annotated in the processed data provided by the authors from the source publication^58^. Analyses were performed using R (v4.2.1/v4.3.1) and the Seurat package (v4.4.0). Data were normalized using the SCTransform() function, and differentially expressed genes (DEGs) were identified with the FindMarkers() function (Wilcoxon rank-sum test). Gene set expression scores were computed using the AddModuleScore() function. UMAP visualizations were generated using DimPlot(), and pie charts, volcano plots, box plots, and violin plots were created with ggplot2 (v3.4.4).

To map single cells to gene expression identities from the CCLE bulk data model, we employed the nnls package (v1.5). Identity enrichment and depletion analyses were performed using GSEA (clusterProfiler v4.6.2) and visualized with enrichplot (v1.18.4). To assess identity expansion or shrinkage, we calculated cell counts for each identity pre- and post-treatment, performing chi-squared tests using chisq.test(). Residual scores from these tests quantified the magnitude of expansion or shrinkage. For cancer cell clusters contributing to identity changes, chi-squared tests were used to assess cluster-specific contributions to the most expanded identity.

#### Single-Cell Multiome Analysis

Raw FASTQ files from single-cell multiome sequencing were aligned to the GRCh38 genome using Cell Ranger ARC (v2.0.1). Downstream analysis was performed in R (v4.2.1/v4.3.1) with the Seurat (v4.4.0) and Signac (v1.10.0) packages to process RNA and ATAC data simultaneously. Datasets for 3 treatment conditions were merged together following the standard pipeline outlined in the Signac tutorial. To ensure high-quality data, single-cell barcodes meeting specific thresholds were retained, including ATAC read counts between 1,000 and 70,000, RNA read counts between 1,000 and 25,000, detection of more than 500 genes per cell, nucleosome signal less than 2, TSS.enrichment more than 1, and mitochondrial transcript proportions below 20%. Doublets were identified and removed using DoubletFinder (v2.0.4).

To validate that all cells within the organoid samples were cancerous, infercnv (v1.21.0), SingleR (v2.4.1) and celldex (v1.12.0) were employed. Reference cells, including T cells, NK cells, macrophages, monocytes, and B cells, from the GEO study GSE176078 were used for infercnv, ensuring that non-cancerous cells were excluded from further analysis. RNA data were normalized using the SCTransform() function in Seurat, with results stored in the “SCT” assay. For ATAC data, The function CallPeaks() was used for peak calling, and the resulting peaks were saved in the “peaks” assay. Gene activity scores, representing transcriptional activity across promoter and gene body regions, were calculated with GeneActivity() from Signac and stored in the “gene_activity” assay.

Transcription factor analysis included identifying chromatin binding motifs with JASPAR2020 (v0.99.10) and quantifying chromatin accessibility associated with these motifs using ChromVAR (v1.20.2). Differential motif accessibility, gene expression changes, and ATAC peaks were identified using the FindMarkers() function in Seurat for each drug treatment condition. Visualization of the data included violin plots created with the VlnPlot() function, while heatmaps, scatter plots, stacked bar plots, and pie charts were generated using ggplot2 (v3.4.4). Chromatin accessibility in genomic regions was visualized with the CoveragePlot() function from Signac.

To systematically identify transcription factors involved in CCLE F3 identity changes after MS177 and JQ1 treatments, upregulated genes—including those in the NF-κB pathway—were input into the Enrichr database (https://maayanlab.cloud/Enrichr/). Potential transcription factors were identified under the “Transcription” section of the results. Gene targets of transcription factors were curated from ChIP-seq databases such as ChEA 2022, ENCODE, and ChEA Consensus TFs from ChIP-X. The intersection of these targets with genes in the MsigDB TNFA signaling hallmark gene set was taken for further analysis.

To evaluate the relationship between transcription factor motif chromatin accessibility and target gene expression, linear models were constructed using the lm() function in R. R-squared values were extracted to quantify the correlation between TF binding motif accessibility and the expression of NF-κB pathway target genes. Differences in R-squared values between treated (JQ1 or MS177) and untreated conditions were calculated and visualized in scatterplots. This analysis highlighted transcription factors whose chromatin accessibility and target gene expression were most affected by drug treatments, revealing molecular drivers of adaptive responses.

For the differential ATAC peaks after drug treatment in the organoid as a whole and in each of the two distinct clusters within the organoid, to link ATAC peaks with genes, the package ChIPseeker(v1.34.1) was employed. Genomic positions for mRNA transcripts were acquired from package TxDb.Hsapiens.UCSC.hg38.knownGene (v3.16.0). Human genome annotation data was acquired from package org.Hs.eg.db (v3.16.0). GRanges objects were created using the package GenomicRange (v1.50.2). The results are stored in **Supplementary Table 8-10** and **Supplementary Table 17-19**.

This integrated analysis pipeline provided a comprehensive view of transcriptional and chromatin accessibility landscapes, enabling the identification of key transcriptional regulators and adaptive mechanisms in TNBC organoid models treated with epigenetic drugs.

## Acknowledgements

The authors acknowledge This work was supported in part by the National Institutes of Health grants K08CA280388, R37CA292075, R01CA280482, P30CA016086 and the UNC Lineberger Center for Triple Negative Breast Cancer

## Author Contributions

Conceptualization, E.B. and P.S.; methodology, E.B., and Y.W.; formal Analysis, Y.W., S.H., and E.B.; funding Acquisition, E.B. and P.S.; investigation, E.B. and P.S.; resources, E.B. and P.S.; supervision, E.B. and P.S.; validation, Y.W.; visualization, Y.W. and E.B.; writing—original draft, E.B. and Y.W.; writing—review and editing, All authors. All authors have read and agreed to the published version of the manuscript.

## Conflicts of Interest

The authors declare no conflict of interest. The funders had no role in the design of the study; in the collection, analyses, or interpretation of data; in the writing of the manuscript, or in the decision to publish the results.

## Notes

### Competing Interest Statement

The authors have declared no competing interest.

