## Extended Data Fig1-10 for "A Multimodal Framework to Uncover Drug-Responsive Subpopulations in Triple-Negative Breast Cancer"

Triple Negative Breast Cancer, TNBC, cell-to-cell heterogeneity, single cell RNA-seq, single cell ATAC-seq, organoid, single cell multiomics, computational genomics, drug response

### Extended Data Figures

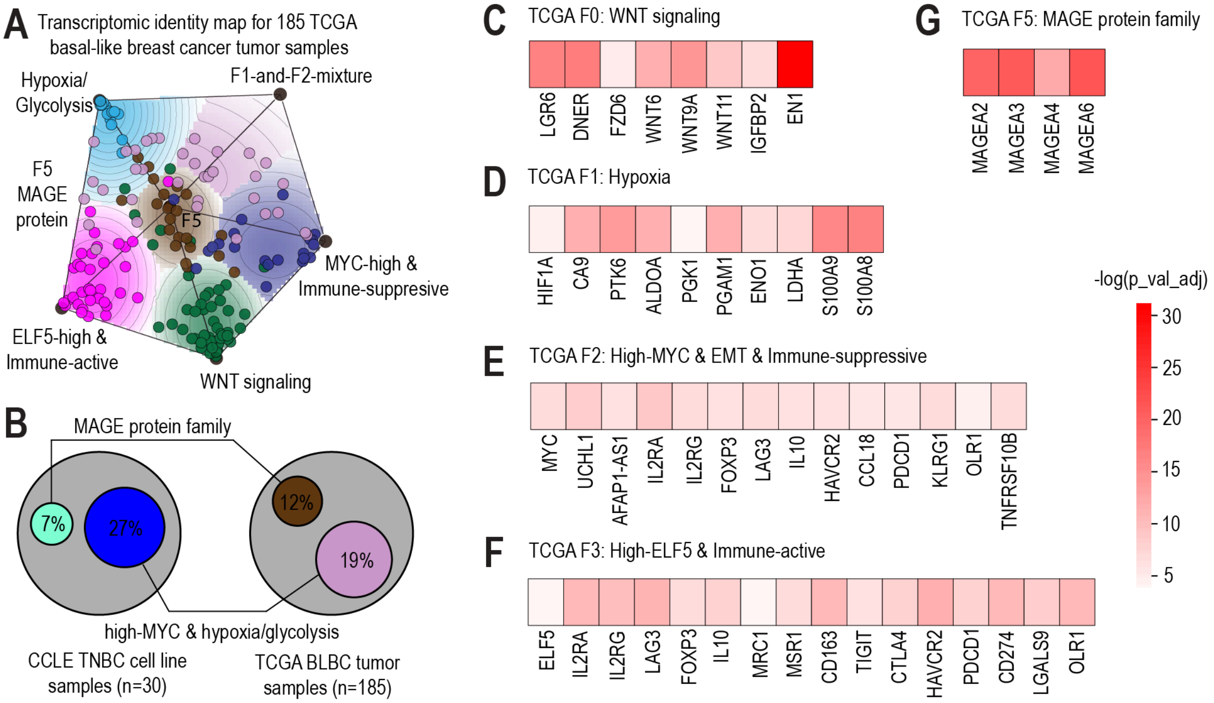


**Extended Data Fig 1**. **TCGA sample identity characterization 1A**. The identity map for 185 TCGA basal-like breast cancer samples. **1B**. Common gene expression identities between cell line and tumor sample models. **1C-1E & 1G**. Gene expression evidence supporting the biology of identities F0, F1, F2 and F5. The heatmaps present the adjusted p-values from Wilcoxon rank-sum tests comparing gene expression levels in tumor samples belonging to a specific identity versus all others. Genes representative of each identity show significantly higher expressions within the associated tumor samples. **1F.** Gene expression evidence supporting the biology of the F3 identity. The heatmap presents the adjusted p-values from Wilcoxon rank-sum tests comparing gene expression levels in tumor samples belonging to the F3 identity versus all others. Gene ELF5 has higher expression in samples belonging to the F3 identity. The rest of genes, which are immune-suppressive genes, have lower expressions in samples belonging to the F3 identity.

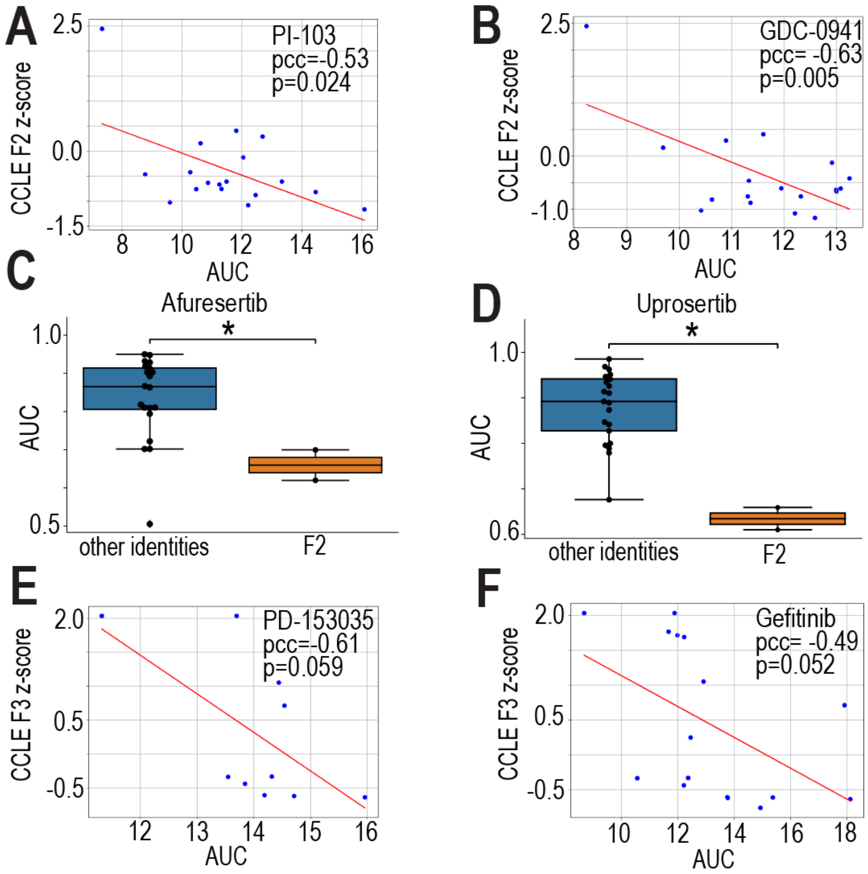


**Extended Data Fig 2. Extra evidence linking drug sensitivity to gene expression identities across TNBC cell lines** Drug efficacies for two PI3K inhibitors PI-103 (**2A**) and GDC-0941 (**2B**) are positively correlated with the CCLE F2 identity z-score across TNBC cell lines. The Pearson correlation coefficient and p-value are shown in the figures. Cell lines belonging to F2 identity are more sensitive to afuresertib (**2C**) and uprosertib (**2D**), two Akt inhibitors. Drug efficacies for two EGFR inhibitors PD-153035 (**2E**) and gefitinib (**2F**) are positively correlated with CCLE F3 identity z-score across TNBC cell lines. The Pearson correlation coefficient and p-value are shown in the figures.

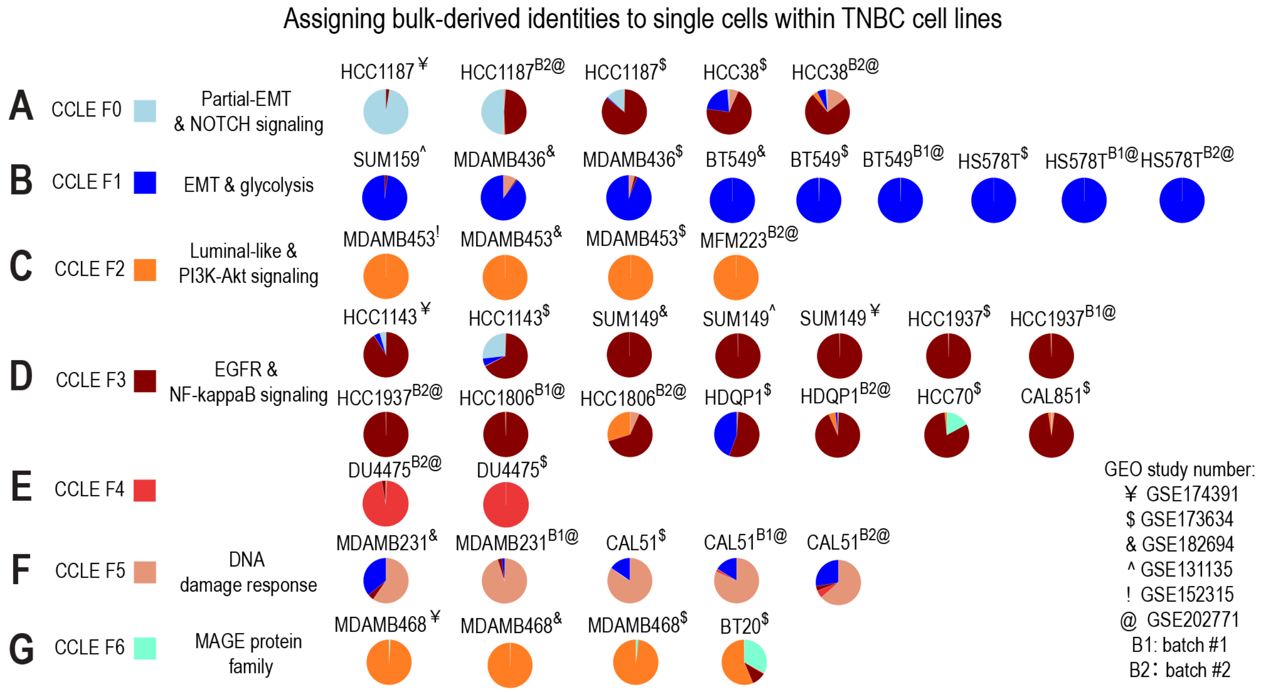


**Extended Data Fig 3. The result of mapping single cells in TNBC cell lines to bulk-derived identities** Each pie chart shows the percentage of single cells belonging to each bulk-level identity in a specific TNBC cell line scRNA-seq dataset. The markers on the upper-right corner of cell line names indicate the GEO study number that the scRNA-seq data is from. Cell line scRNA-seq data is organized based on the bulk-identity it belongs to. The mapping result of single cells in TNBC cell lines from CCLE F0-F6 identities are shown in **3A-3G** correspondingly.

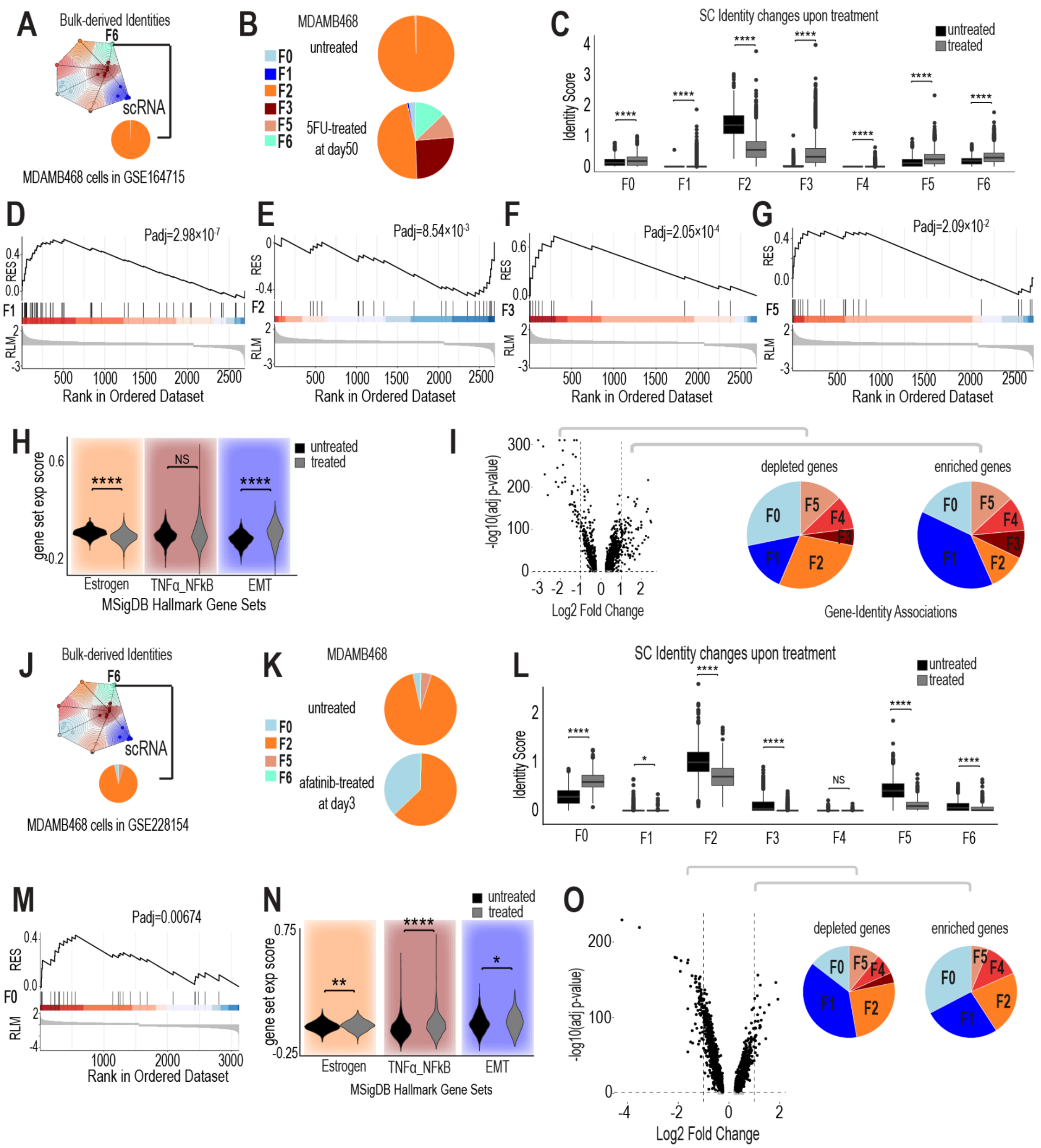


**Extended Data Fig 4. Single-cell identity changes in MDAMB468 after treatments by 5-FU and afatinib 4A.** Single-cell identity composition of MDAMB468 from GSE164715, a TNBC cell line associated with the F6 identity in the CCLE model. **4B**. Single-cell identity shifts in MDAMB468 from GSE164715 after 5-FU treatment. **4C**. Identity scores for single cells in MDAMB468 from GSE164715 before and after 5-FU treatment. Gene Set Enrichment Analysis (GSEA) results for F1 (**4D**), F2 (**4E**), F3 (**4F**), and F5 (**4G**) gene sets after 5-FU treatment. **4H**. Expression scores for three MsigDB hallmark gene sets in MDAMB468 single cells from GSE164715 before and after 5-FU treatment. **4I.** Volcano plots of differentially expressed genes

(DEGs) in MDAMB468 single cells from GSE164715 after 5-FU treatment. For upregulated genes, the pie chart on the right illustrates the proportion of DEGs belonging to each identity as defined in the CCLE model. Similarly, for downregulated genes, the pie chart on the left shows the identity distribution of DEGs. **4J.** Single-cell identity composition of MDAMB468 from GSE228154, a TNBC cell line associated with the F6 identity in the CCLE model. **4K**. Single-cell identity shifts in MDAMB468 from GSE228154 after afatinib treatment. **4L**. Identity scores for MDAMB468 single cells from GSE228154 before and after afatinib treatment. Gene Set Enrichment Analysis (GSEA) results for the F0 (**4M**) gene set after afatinib treatment. **4N**. Expression scores for three MsigDB hallmark gene sets in MDAMB468 single cells from GSE228154 before and after afatinib treatment. **4O.** Volcano plots of differentially expressed genes (DEGs) in MDAMB468 single cells after afatinib treatment. For upregulated genes, the pie chart on the right illustrates the proportion of DEGs belonging to each identity as defined in the CCLE model. Similarly, for downregulated genes, the pie chart on the left shows the identity distribution of DEGs.

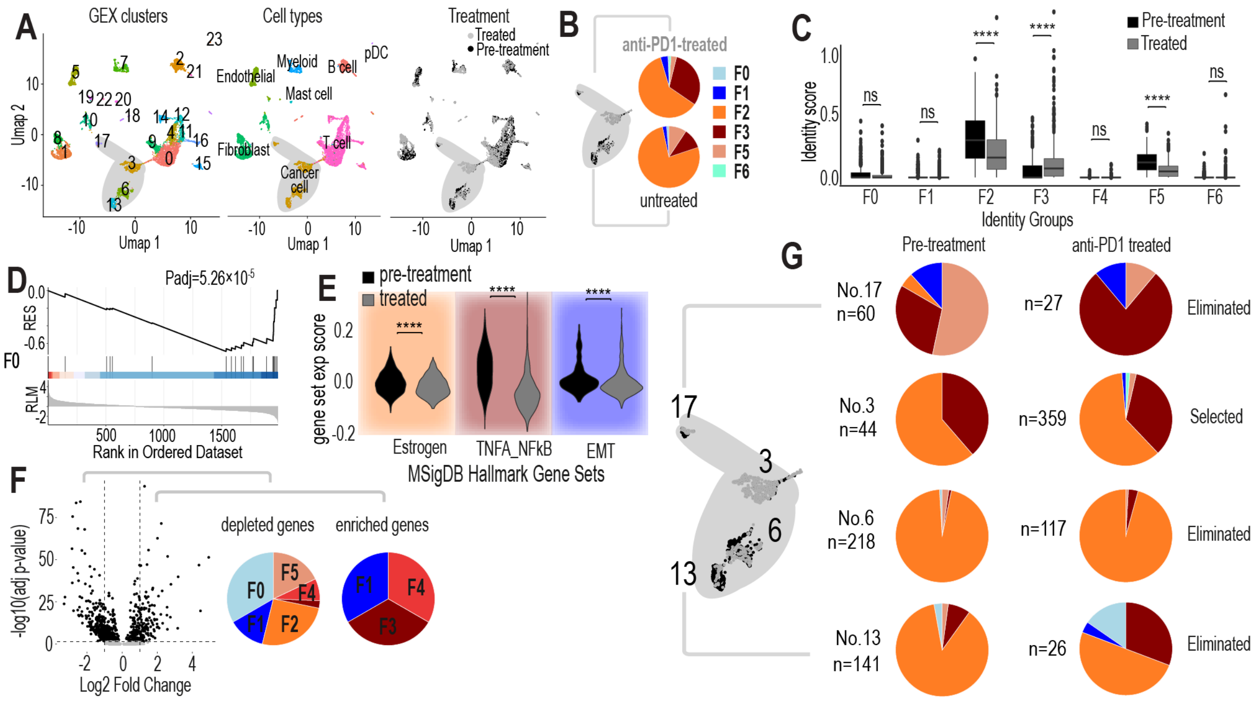


**Extended Data Fig 5. Single-cell identity changes of cancer cells in patient sample BIOKEY_19 after anti-PD1 treatment 5A.** UMAP plots showing cluster identifications, cell types, and treatment conditions for all single cells in the BIOKEY_19 tumor. These plots provide an overview of the cellular landscape within the tumor, highlighting distinct clusters and their associations with treatment conditions. **5B.** Single-cell identity changes in the cancer cells of TNBC patient BIOKEY_19 after pembrolizumab treatment. Pie charts illustrate the distribution of single cells across different identities within the cancer cell population before and after treatment. **5C.** Identity scores for single cells in the BIOKEY_19 cancer cell population before and after pembrolizumab treatment. **5D.** GSEA results for the F0 gene set after pembrolizumab treatment, highlighting the enrichment or depletion of these identity-specific gene sets in the treated cancer cell population. **5E.** Expression scores for three MsigDB hallmark gene sets in

BIOKEY_19 cancer single cells before and after treatment. The hallmark gene sets analyzed include estrogen response (early and late combined), TNFα signaling via NF-κB, and epithelial-mesenchymal transition (EMT), providing insights into treatment-induced changes in key biological pathways. **5F.** Volcano plots of differentially expressed genes (DEGs) in BIOKEY_19 cancer single cells after pembrolizumab treatment. Pie charts accompanying the plots display the percentage of DEGs associated with each identity in the CCLE model. The right pie chart represents upregulated genes, while the left represents downregulated genes, linking these changes to specific identities. **5G.** Analysis of identity composition changes in four cancer cell clusters from BIOKEY_19 (cluster_3, cluster_6, cluster_13, and cluster_17) before and after pembrolizumab treatment. Pie charts illustrate the percentage of single cells belonging to each identity within each cluster, revealing differential responses across clusters.

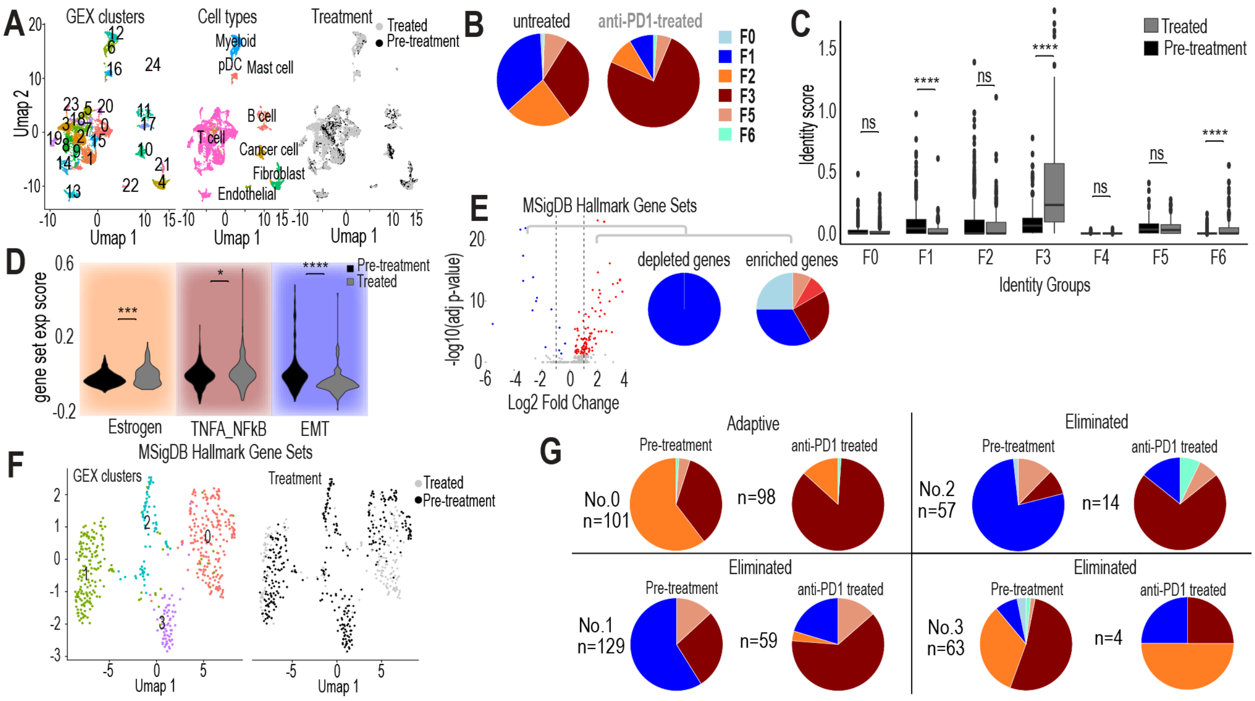


**Extended Data Fig 6. Single-cell identity changes of cancer cells in patient sample BIOKEY_10 after anti-PD1 treatment 6A.** UMAP plots showing cluster identifications, cell types, and treatment conditions for all single cells in the BIOKEY_10 tumor. These plots provide an overview of the cellular landscape within the tumor, highlighting distinct clusters and their associations with treatment conditions. **6B.** Single-cell identity changes in the cancer cells of TNBC patient BIOKEY_10 after pembrolizumab treatment. Pie charts illustrate the distribution of single cells across different identities within the cancer cell population before and after treatment. **6C.** Identity scores for single cells in the BIOKEY_10 cancer cell population before and after pembrolizumab treatment. **6D.** Expression scores for three MsigDB hallmark gene sets in BIOKEY_10 cancer single cells before and after treatment. The hallmark gene sets analyzed include estrogen response (early and late combined), TNFα signaling via NF-κB, and epithelial-mesenchymal transition (EMT), providing insights into treatment-induced changes in key biological pathways. **6E.** Volcano plots of differentially expressed genes (DEGs) in BIOKEY_10 cancer single cells after pembrolizumab treatment. Pie charts accompanying the plots display the percentage of DEGs associated with each identity in the CCLE model. The right pie chart represents upregulated genes, while the left represents downregulated genes, linking these changes to specific identities. **6F**. UMAP plots showing cluster identifications and treatment conditions for cancer cells in the BIOKEY_10 tumor. **6G.** Analysis of identity composition changes in four cancer cell clusters from **Extended Data Fig 6F** (cluster_0, cluster_1, cluster_2, and cluster_3) before and after pembrolizumab treatment. Pie charts illustrate the percentage of single cells belonging to each identity within each cluster, revealing differential responses across clusters.

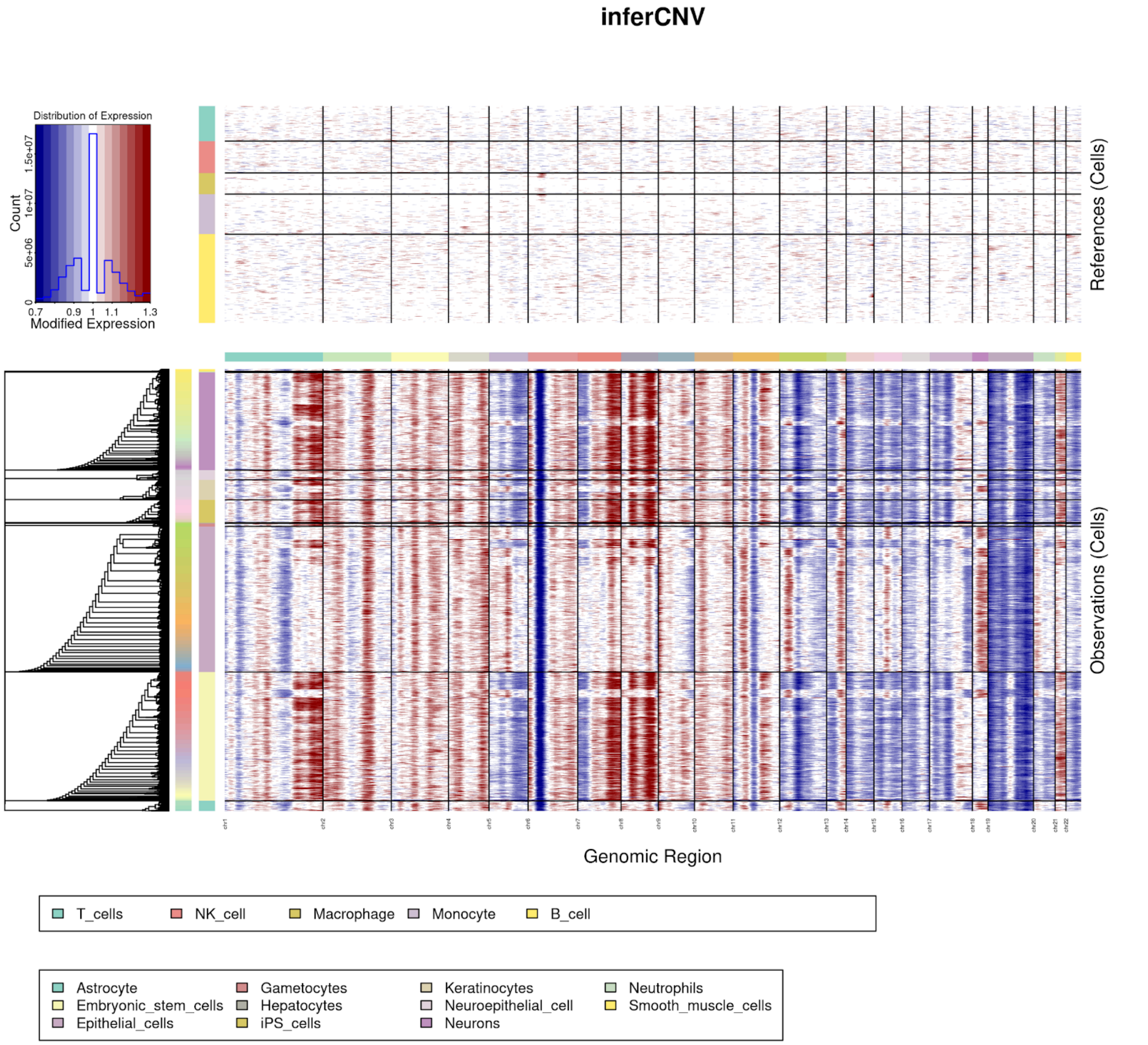


**Extended Data Fig 7. Infercnv result for cells in the organoid** The top heatmap shows the chromosomal copy number alteration profiles in the reference cells. The bottom heatmap shows the chromosomal copy number alteration profiles of the single cells in the organoid. In the heatmap, each row stands for a cell. Each column stands for a gene. Genes are arranged based on their chromosomal locations.

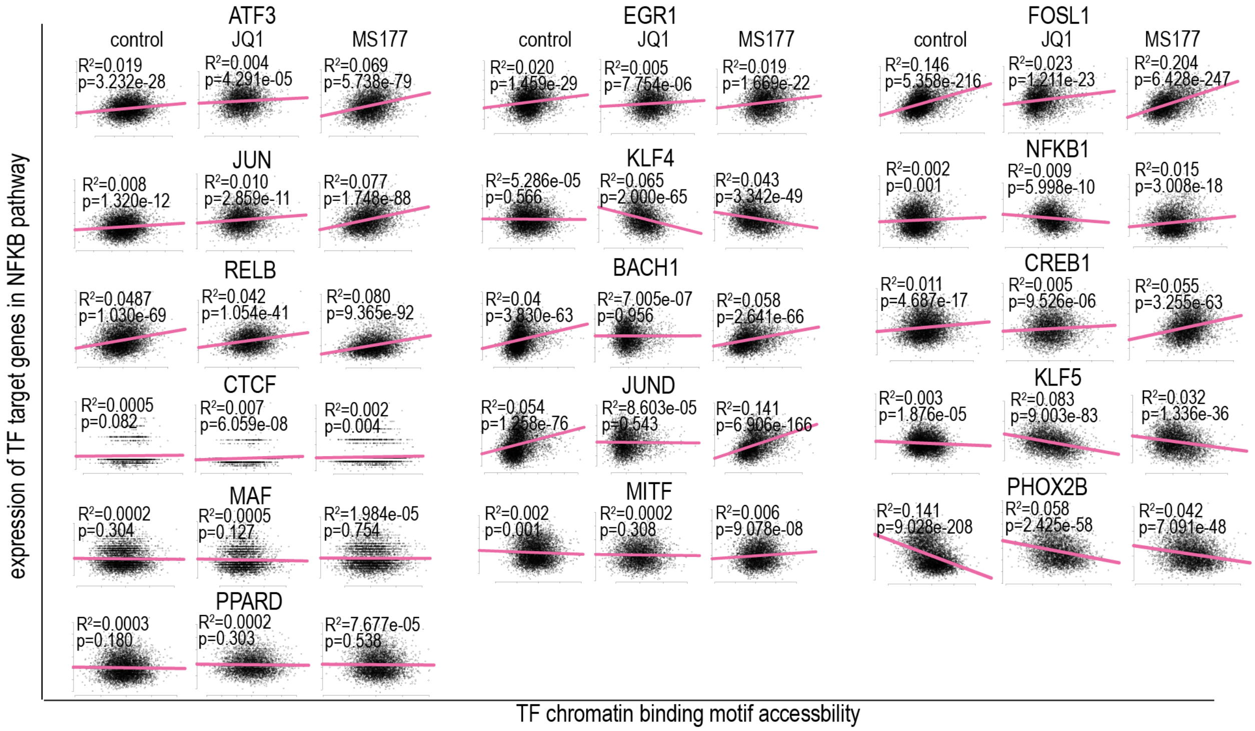


**Extended Data Fig 8. The relationship between TF chromatin binding motif accessibility and the expression of its target genes in NF-κB signaling pathway for all the TFs shown in main Fig. 5I in control, JQ1-treated, and MS177-treated conditions** Comparing all of the TFs shown here with RELA, RELA has the strongest regulation of NF-κB pathway gene expression in the MS177-treated condition, and it is the most involved TF in response to MS177 treatment by comparing $R^2$ difference between MS177 and control conditions.

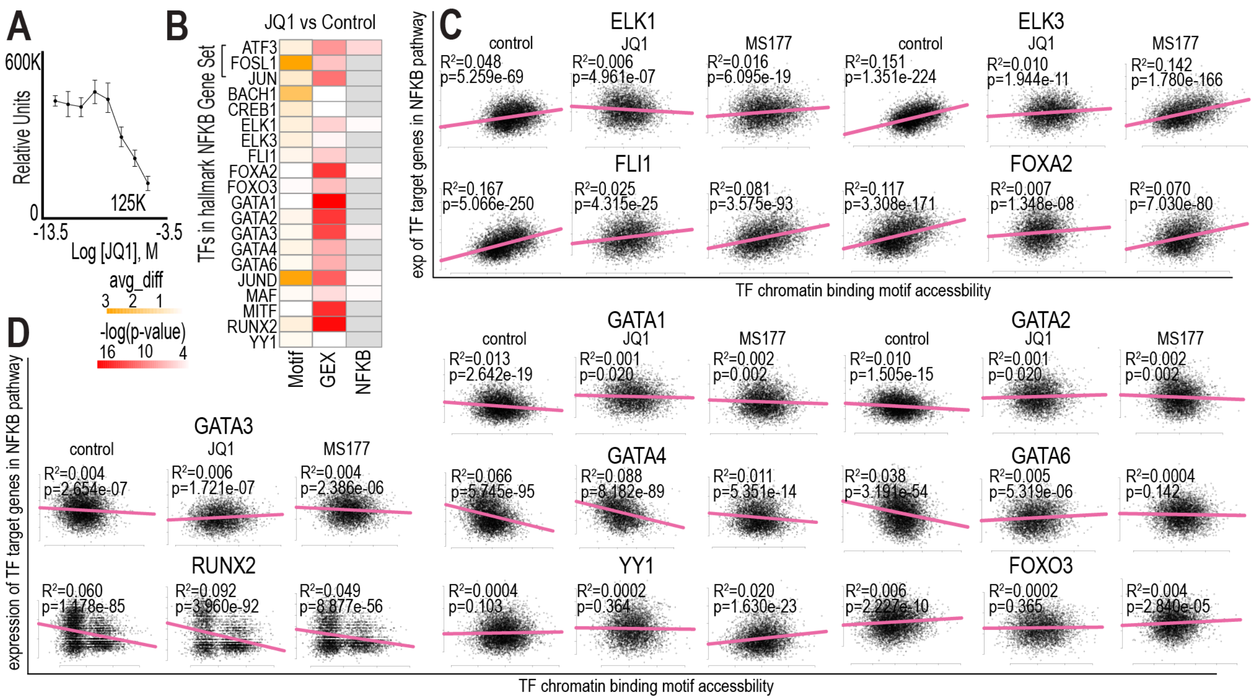


**Extended Data Fig 9. TFs potentially involved in JQ1 drug response. 9A.** Drug treatment curve showing the response of the organoid to JQ1, indicating the relative efficacy of the treatment on the organoid population **9B.** Analysis of transcription factors (TFs) relevant to the response to JQ1 treatment. The first column shows the average difference in TF binding motif accessibility after JQ1 treatment. The second column highlights the significance of TFs in upregulating gene expression broadly after JQ1 treatment. The third column focuses on TFs that significantly upregulate gene expression specifically in NF-κB pathway genes after JQ1 treatment. **9C-9D.** The relationship between TF chromatin binding motif accessibility and the expression of its target genes in NF-κB signaling pathway for all the TFs shown in **Extended Data Fig 9A** in the control, JQ1-treated, and MS177-treated conditions. Comparing all of the TFs shown here with RELA, RELA has the strongest regulation of NF-κB pathway gene expression in the MS177-treated condition, and it is the most involved TF in response to MS177 treatment by comparing $R^2$ difference between MS177 and control conditions.

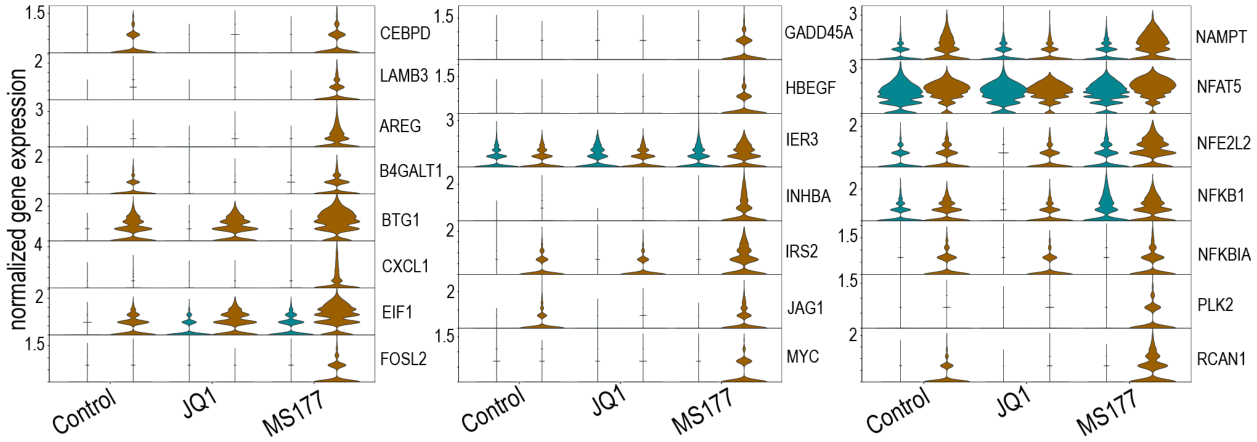


**Extended Data Fig 10. Genes in the NF-κB hallmark gene set whose expressions are specifically induced by MS177 in the stem-cell-like cluster** This further confirms that distinct clusters of cancer cells have different pathway activities in response to MS177 treatment.
