## Supplementary Information Fig 1-11 for "A Multimodal Framework to Uncover Drug-Responsive Subpopulations in Triple-Negative Breast Cancer"

Triple Negative Breast Cancer, TNBC, cell-to-cell heterogeneity, single cell RNA-seq, single cell ATAC-seq, organoid, single cell multiomics, computational genomics, drug response

### Supplementary Information

CCLE F0 Identity---Notch signaling

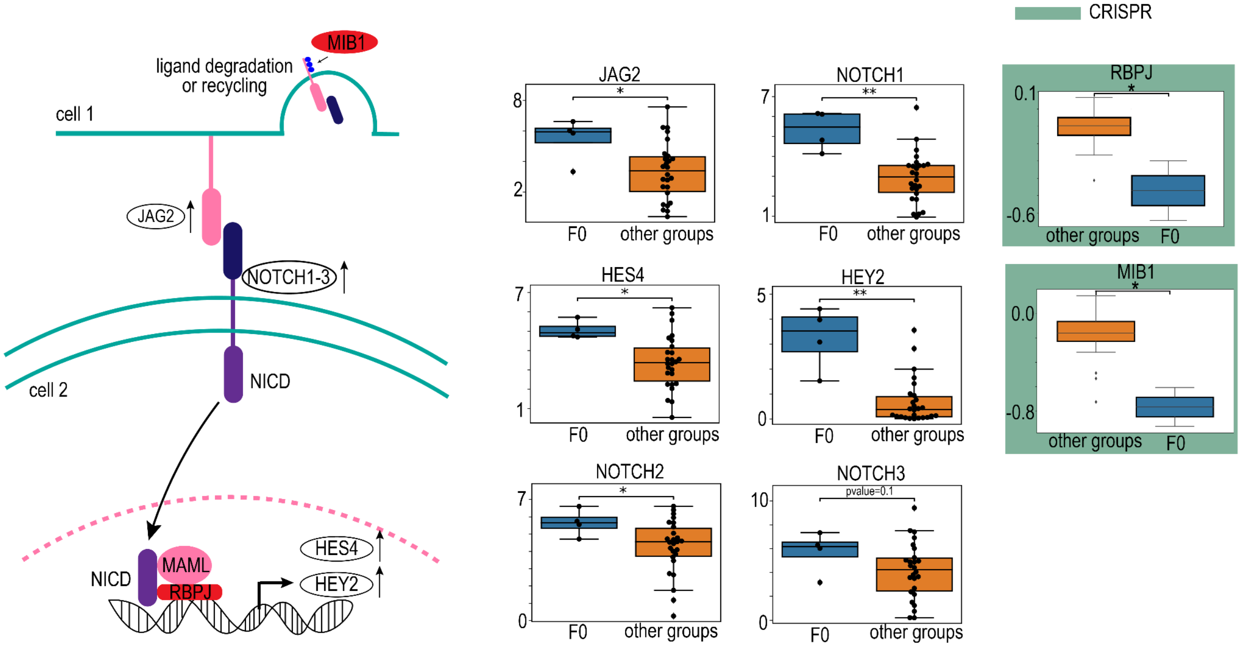


cartoon figure adapted from Zhao F, He Y, Zhao Z, et al. The Notch signaling-regulated angiogenesis in rheumatoid arthritis: pathogenic mechanisms and therapeutic potentials. Front Immunol. 2023;14:1272133. Published 2023 Oct 26. doi:10.3389/fimmu.2023.1272133

**Supplementary Information Fig 1. Cartoon illustration demonstrating the involvement of CCLE F0 identity-defining genes in Notch signaling** NICD: Notch intracellular domain, MAML: mastermind-like proteins

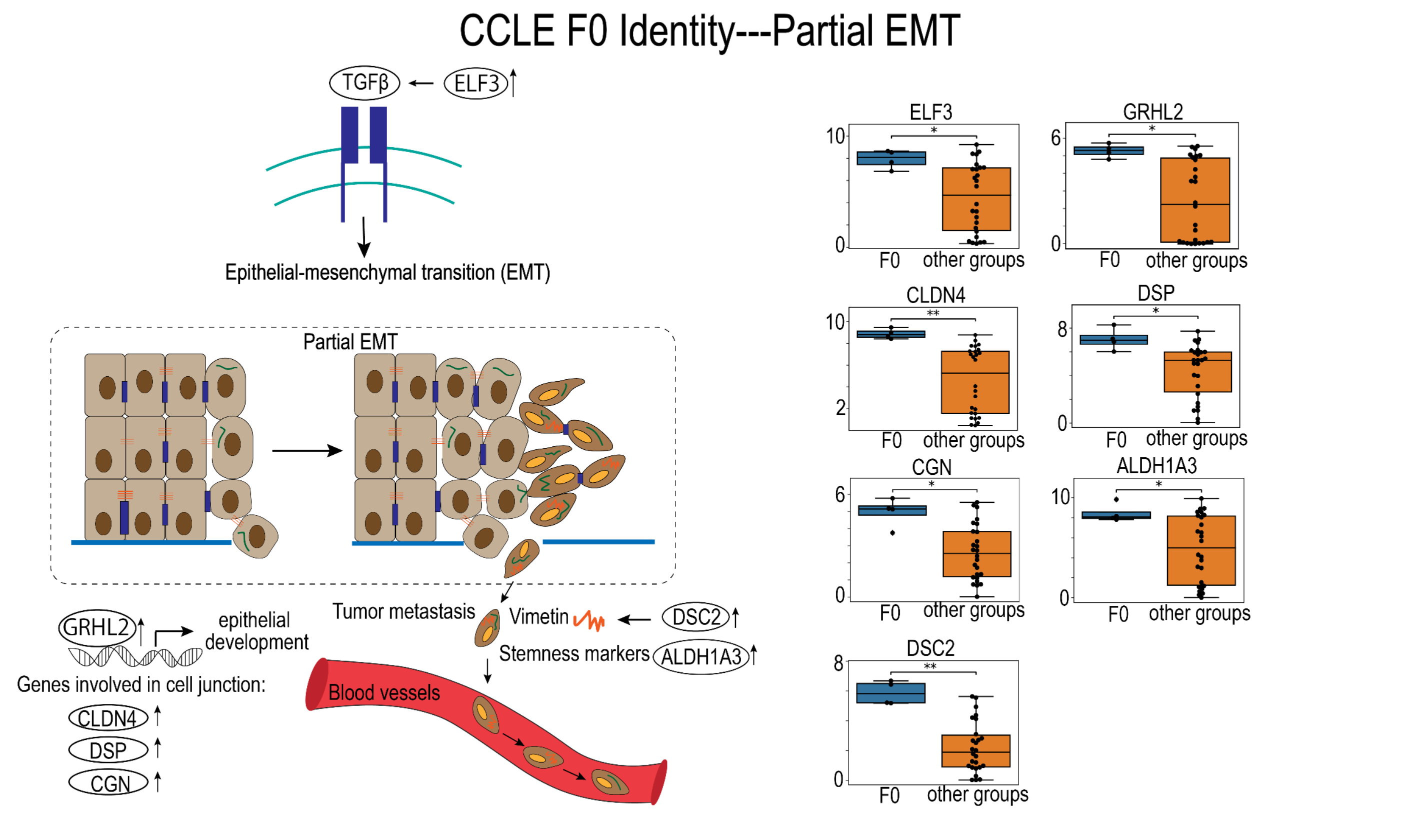


cartoon figure adapted from Sheng L, Zhuang S. New Insights Into the Role and Mechanism of Partial Epithelial-Mesenchymal Transition in Kidney Fibrosis. Front Physiol. 2020;11:569322. Published 2020 Sep 15. doi:10.3389/fphys.2020.569322

**Supplementary Information Fig 2. Cartoon illustration demonstrating the involvement of CCLE F0 identity-defining genes in partial-EMT**

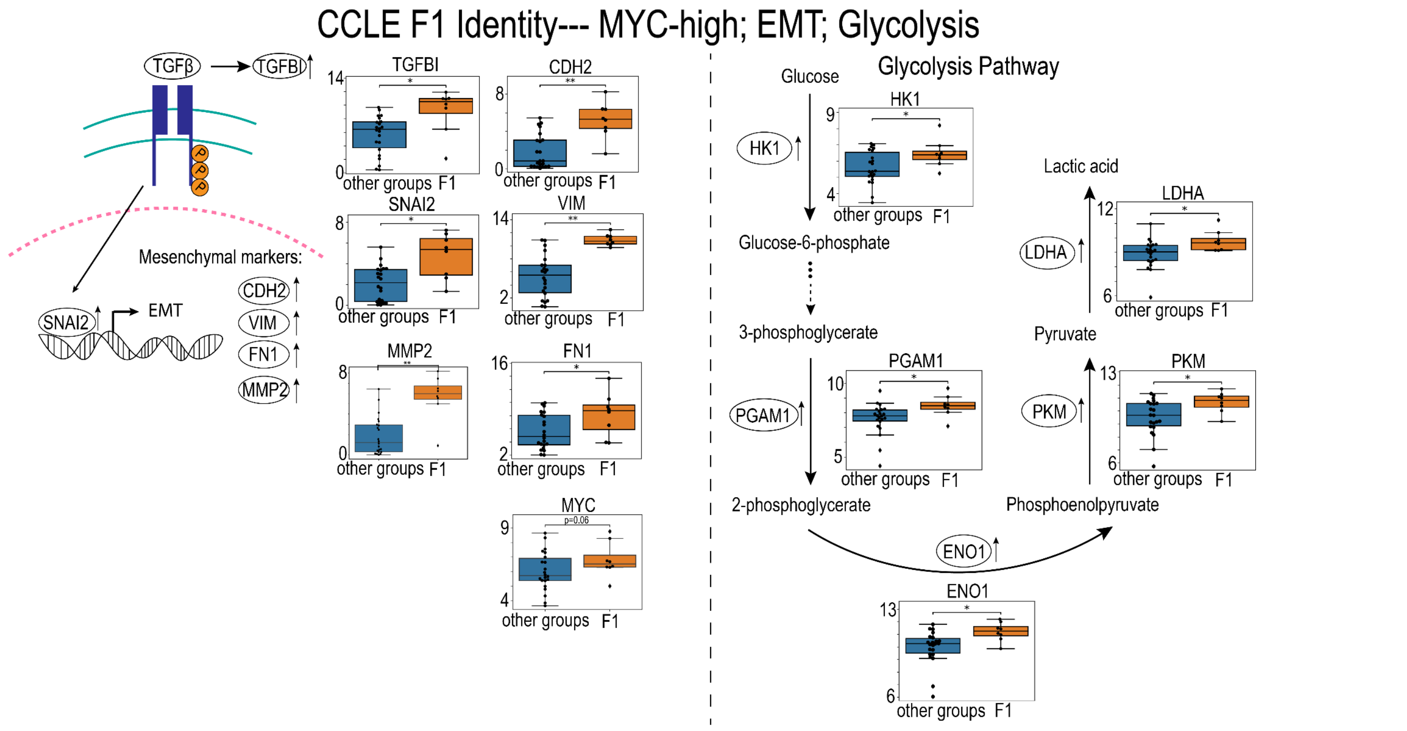

**Supplementary Information Fig 3. Cartoon illustration demonstrating the involvement of CCLE F1 identity-defining genes in MYC-high, EMT and glycolysis**

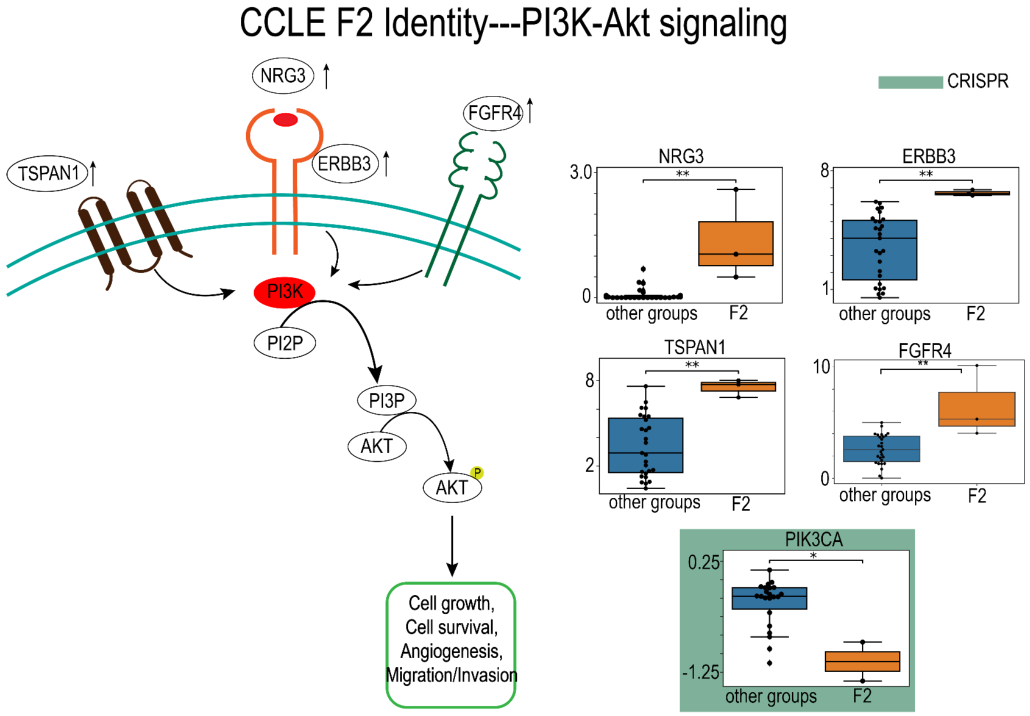


cartoon figure adapted from Garcia-Mayea Y, Mir C, Carballo L, Sánchez-Garcia A, Bataller M, LLeonart ME. TSPAN1, a novel tetraspanin member highly involved in carcinogenesis and chemoresistance. Biochim Biophys Acta Rev Cancer. 2022;1877(1):188674. doi:10.1016/j.bbcan.2021.188674

**Supplementary Information Fig 4. Cartoon illustration demonstrating the involvement of CCLE F2 identity-defining genes in PI3K-Akt signaling**

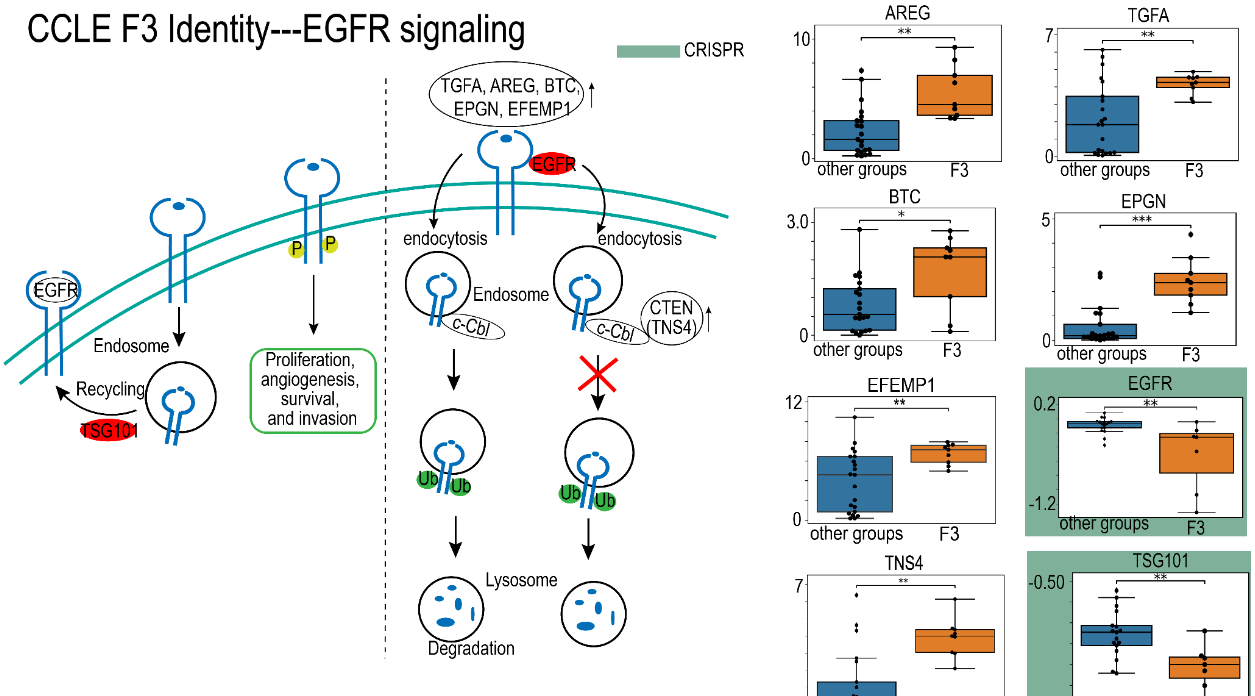


cartoon figure adapted from Xiao D, Hu X, Peng M, et al. Inhibitory role of proguanil on the growth of bladder cancer via enhancing EGFR degradation and inhibiting its downstream signaling pathway to induce autophagy. Cell Death Dis. 2022;13(5):499. Published 2022 May 25. doi:10.1038/s41419-022-04937-z

**Supplementary Information Fig 5. Cartoon illustration demonstrating the involvement of CCLE F3 identity-defining genes in EGFR signaling.** c-Cbl: Casitas B lineage lymphoma, an E3 ubiquitin ligase

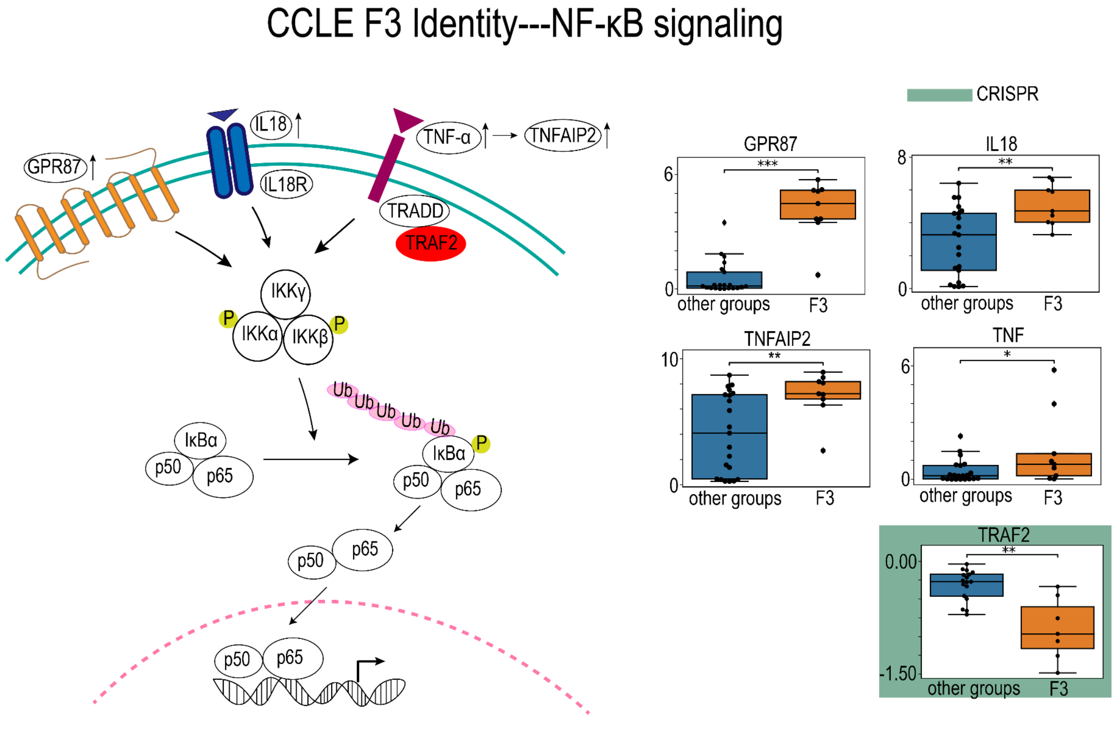


cartoon figure adapted from Peng C, Ouyang Y, Lu N, Li N. The NF-κB Signaling Pathway, the Microbiota, and Gastrointestinal Tumorigenesis: Recent Advances. Front Immunol. 2020;11:1387. Published 2020 Jun 30. doi:10.3389/fimmu.2020.01387

**Supplementary Information Fig 6. Cartoon illustration demonstrating the involvement of CCLE F3 identity-defining genes in NF-κB signaling**

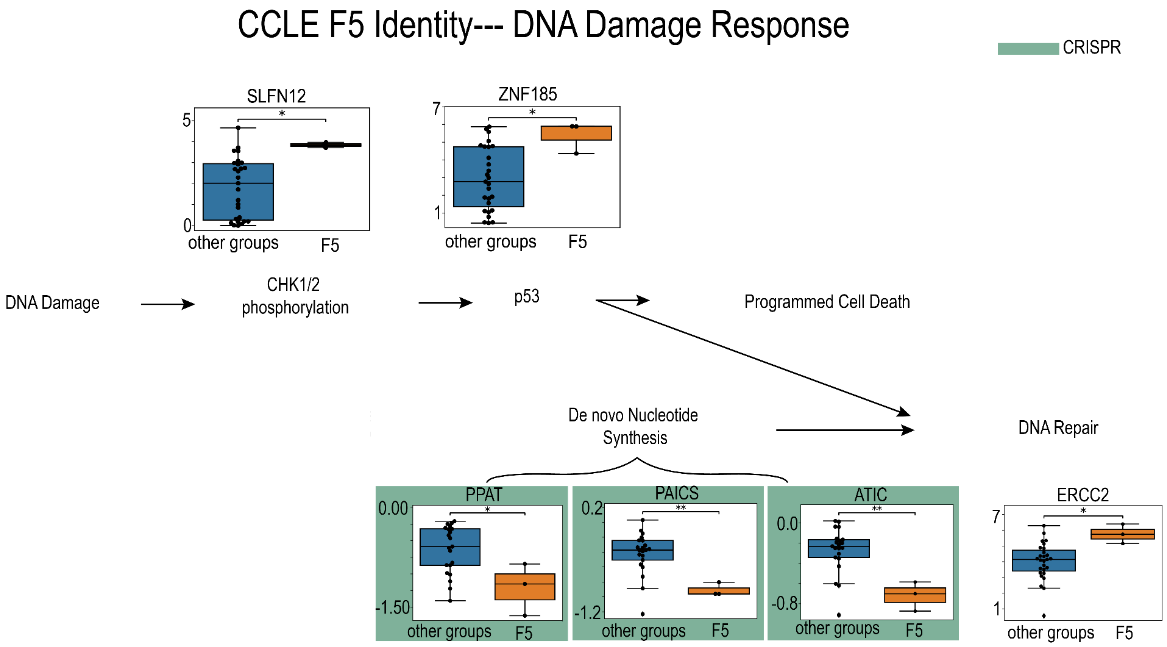


**Supplementary Information Fig 7. Cartoon illustration demonstrating how identity-defining genes point to DNA damage response in CCLE F5 identity**

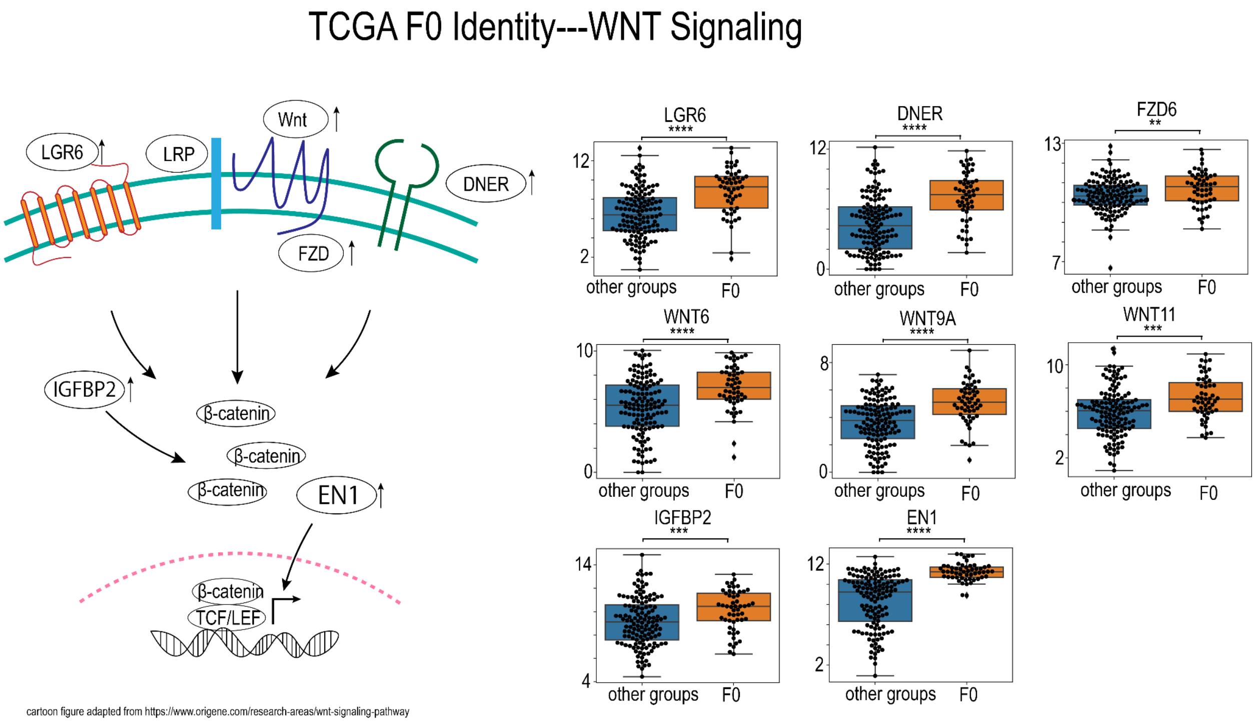

**Supplementary Information Fig 8. Cartoon illustration demonstrating how identity-defining genes point to WNT signaling in TCGA F0 identity** LRP: low-density lipoprotein receptor-related protein, TCF/LEF: T-cell factor/lymphoid enhancer factor

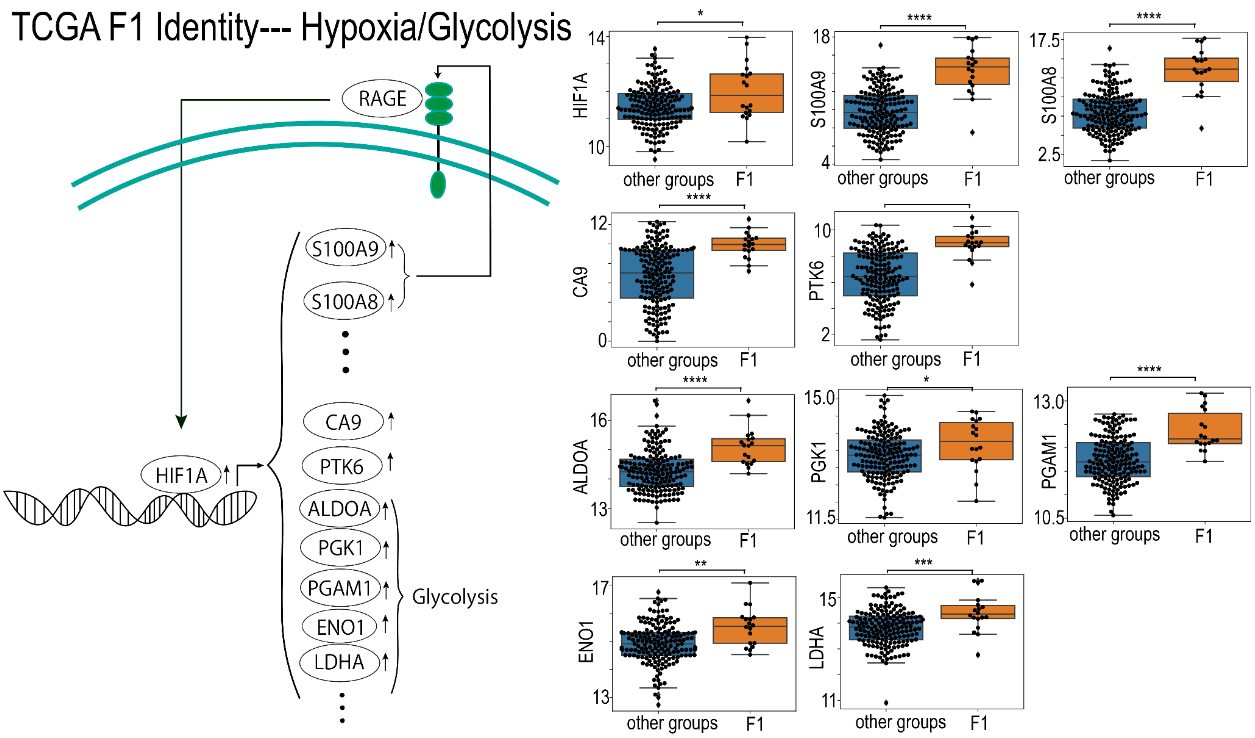


**Supplementary Information Fig 9. Cartoon illustration demonstrating how identity-defining genes point to hypoxia/glycolysis in TCGA F1 identity.** RAGE: receptor for advanced glycation end products

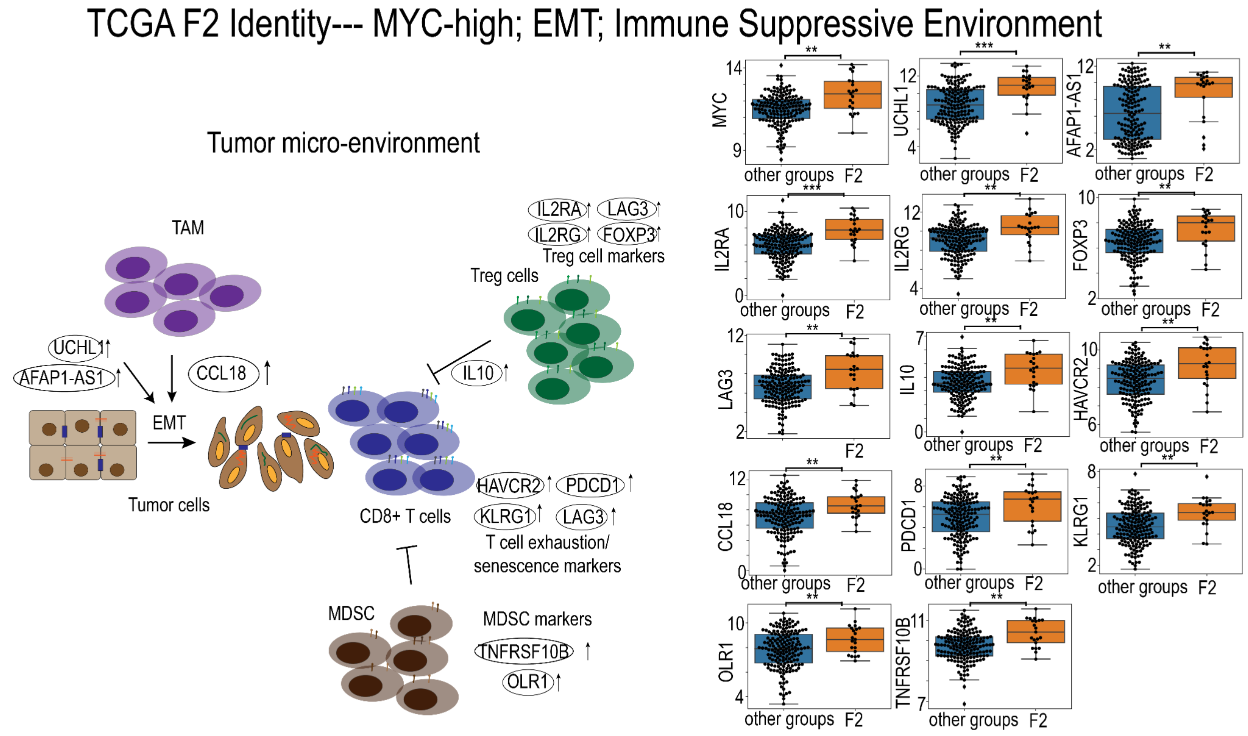


**Supplementary Information Fig 10. Cartoon illustration demonstrating the involvement of TCGA F2 identity-defining genes in "MYC"-high, EMT and immune suppressive environment.** TAM: tumor-associated macrophage, MDSC: myeloid-derived suppressor cell

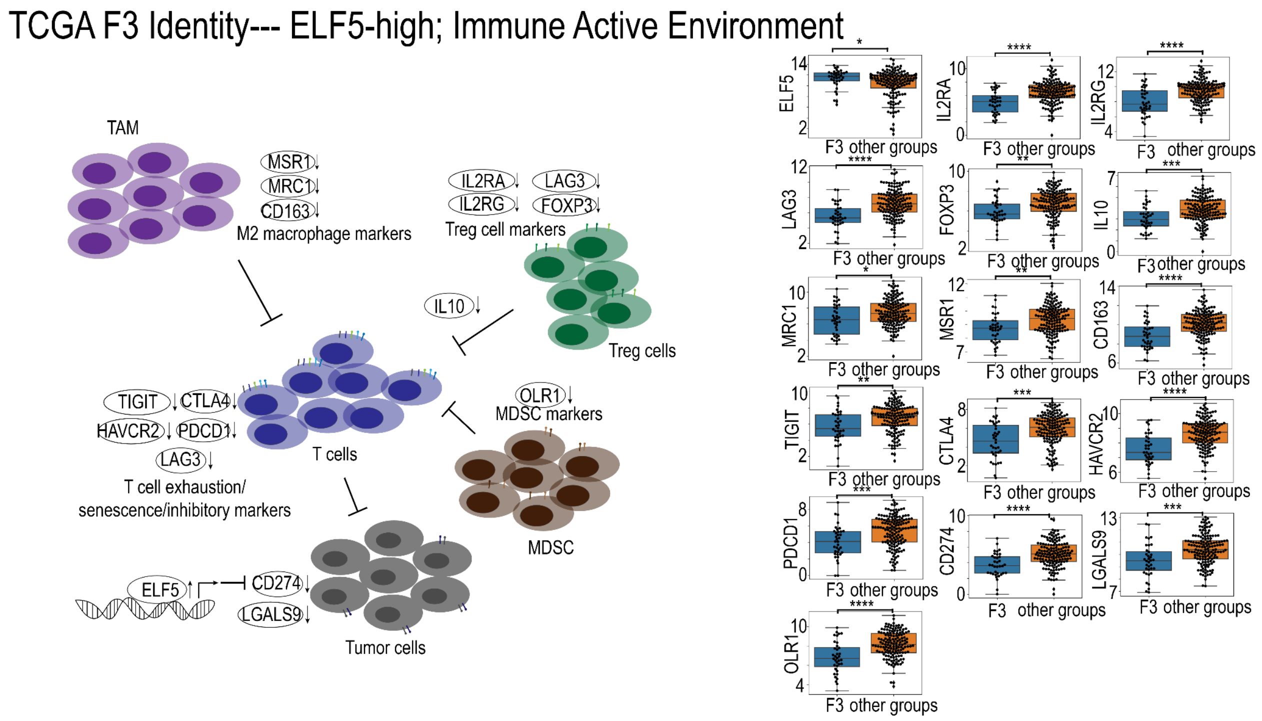

**Supplementary Information Fig 11. Cartoon illustration demonstrating the involvement of TCGA F3 identity-defining genes in "ELF5"-high and immune active environment** TAM: tumor-associated macrophage, MDSC: myeloid-derived suppressor cell
